# Hemodynamic timing in resting-state and breathing-task BOLD fMRI

**DOI:** 10.1101/2022.11.11.516194

**Authors:** Jingxuan Gong, Rachael C. Stickland, Molly G. Bright

## Abstract

The blood flow response to a vasoactive stimulus demonstrates regional heterogeneity across both the healthy brain and in cerebrovascular pathology. The timing of a regional hemodynamic response is emerging as an important biomarker of cerebrovascular dysfunction, as well as a confound within fMRI analyses. Previous research demonstrated that hemodynamic timing is more robustly characterized when a larger systemic vascular response is evoked by a breathing challenge, compared to when only spontaneous fluctuations in vascular physiology are present (i.e., in resting-state data). However, it is not clear whether hemodynamic delays in these two conditions are physiologically interchangeable, and how methodological signal-to-noise factors may limit their agreement. To address this, we generated whole-brain maps of hemodynamic delays in nine healthy adults. We assessed the agreement of voxel-wise gray matter (GM) hemodynamic delays between two conditions: resting-state and breath-holding. We found that delay values demonstrated poor agreement when considering all GM voxels, but increasingly greater agreement when limiting analyses to voxels showing strong correlation with the GM mean time-series. Voxels showing the strongest agreement with the GM mean time-series were primarily located near large venous vessels, however these voxels explain some, but not all, of the observed agreement in timing. Increasing the degree of spatial smoothing of the fMRI data enhanced the correlation between individual voxel time-series and the GM mean time-series. These results suggest that signal-to-noise factors may be limiting the accuracy of voxel-wise timing estimates and hence their agreement between the two data segments. In conclusion, caution must be taken when using voxel-wise delay estimates from resting-state and breathing-task data interchangeably, and additional work is needed to evaluate their relative sensitivity and specificity to aspects of vascular physiology and pathology.

## 1. Introduction

Oxygenated blood takes time to arrive at the brain, traveling from the heart and through the vascular tree to ultimately perfuse into local brain tissue where nutrient exchange can occur. Changes in blood flow can lead to changes in blood oxygenation, particularly in the venous vessels downstream of the exchange with tissue. Functional Magnetic Resonance Imaging (fMRI) is sensitive to blood flow changes due to their impact on local blood oxygenation levels, referred to as Blood Oxygenation Level Dependent (BOLD) contrast (Glover, 2011). However, processes which initiate local blood flow changes will not produce instantaneous changes in BOLD fMRI signals. A single neural event evokes a local hemodynamic response through neurovascular coupling mechanisms (Huneau, Benali, & Chabriat, 2015), and this response is typically modeled with the canonical hemodynamic response function or HRF (Friston et al., 1998; Martindale et al., 2003; Penny, Friston, Ashburner, Kiebel, & Nichols, 2006). Determined empirically, this HRF peaks many seconds after an impulse of neural stimulation suggesting that there is an inherent BOLD fMRI “delay” due to the evolution of the local blood flow response to nearby neural activity. There is evidence that the shape and timing of the HRF to a neural event may vary across brain regions (Handwerker, Ollinger, & D’Esposito, 2004), with healthy aging (Issard & Gervain, 2018; West et al., 2019), and under different baseline states (Cohen, Ugurbil, & Kim, 2002; Liu et al., 2004). Via a different process to neurovascular coupling, blood flow changes can be evoked systemically by a non-neural vasodilatory stimulus such as carbon dioxide, a phenomenon known as cerebrovascular reactivity (CVR) (Chen & Gauthier, 2021; Liu, De Vis, & Lu, 2019; Pinto, Bright, Bulte, & Figueiredo, 2020; Sleight, Stringer, Marshall, Wardlaw, & Thrippleton, 2021). These CVR responses are also delayed, with regional heterogeneity in the vascular response observed across the healthy brain and altered by disease (Bright, Bulte, Jezzard, & Duyn, 2009; Donahue et al., 2016; Holmes et al., 2020; Leung, Duffin, Fisher, & Kassner, 2016; Moia et al., 2020; Siegel, Snyder, Ramsey, Shulman, & Corbetta, 2016; Sousa, Vilela, & Figueiredo, 2014). There are several additional non-neural phenomena thought to drive variability in the spontaneous and systemic low frequency oscillation (sLFO) in BOLD contrast. Typically defined below 0.15Hz, evidence of sources of variation in this non-stationary sLFO point towards blood CO_2_, heart rate and respiratory volume variations, gastric oscillations, vasomotion (Tong, Hocke, & Frederick, 2019), and vascular structure (Aso, Jiang, Urayama, & Fukuyama, 2017) including specifically deoxyhemoglobin density (Aso, Urayama, Fukuyama, & Murai, 2019).

With growing evidence that hemodynamic timing is a useful clinical metric in pathologies such as moyamoya disease (Donahue et al., 2016; Jahanian, Christen, Moseley, & Zaharchuk, 2018), carotid stenosis (Chang et al., 2013), stroke (Ni et al., 2017; Siegel et al., 2016), glioma (Cai et al., 2022), and Alzheimer’s (Holmes et al., 2020), we must be cautious when interpreting fMRI timing information that is derived from different stimuli or conditions. In some experimental circumstances, it may be desirable to deliver a controlled and substantial vasodilatory stimulus to enhance the effect size of the corresponding BOLD signal changes and make characterization of hemodynamic timings more robust. Researchers use complex gas inhalation challenges (Liu et al., 2019) as well as simple breathing tasks (e.g., breath-holding) in order to modulate blood CO_2_ levels and evoke a large systemic vascular response. Breathing-tasks have been used successfully to map the variability of BOLD hemodynamic timings in both healthy and patient populations (Bright et al., 2009; Chang, Thomason, & Glover, 2008; Geranmayeh, Wise, Leech, & Murphy, 2015; Magon et al., 2009; Moia et al., 2021; Pinto, Jorge, Sousa, Vilela, & Figueiredo, 2016; Raut, Nair, Sattin, & Prabhakaran, 2016; Sousa et al., 2014). We have previously demonstrated that the timing of the BOLD CVR response is more robustly characterized when a large systemic vascular response is evoked by a breathing challenge, compared to when only spontaneous fluctuations in arterial CO_2_ levels are present (Bright & Murphy, 2017; Stickland et al., 2021). Still, researchers have had success with mapping BOLD hemodynamic timings in healthy and patient populations using resting-state fMRI data in the absence of evoked responses (Cai et al., 2022; Hu et al., 2021; Liu, Li, et al., 2017; Siegel et al., 2016; Tong et al., 2019). Despite the anticipated lower signal-to-noise ratio of the vascular effects, this resting-state experimental design is simpler and therefore suitable for a wider range of populations and clinical or research contexts.

However, we anticipate that the mechanisms underpinning voxel-wise and global hemodynamics in these two conditions, breathing-task and resting-state fMRI, may differ, making it unclear whether hemodynamic timings derived from these two types of data provide interchangeable physiological information. Although it is reasonable to assume that the majority of information present in the fMRI time-series during a breath-holding task reflects the associated evoked CVR response, there will be some neural activity associated with the voluntary performance of the task, in addition to a baseline level of intrinsic neuronal fluctuations. Similarly, intrinsic low-frequency fluctuations in resting-state fMRI signals are frequently used to characterize neural activation patterns of intrinsic brain networks, but it is well established that many physiologic phenomena, both neural and vascular, influence blood flow and BOLD fMRI signals in uncontrolled ways (Caballero-Gaudes & Reynolds, 2017; Liu, 2013, 2016; Murphy, Birn, & Bandettini, 2013; Tong et al., 2019). Finally, systemic physiological drivers of LFOs other than CO_2_ will impact both conditions (Aso et al., 2019). Thus, these two types of fMRI data – breathing-task and resting-state – contain diverse types of neural and vascular information, albeit with different weightings.

In this paper, we estimate relative hemodynamic timings across the brain in BOLD fMRI data from healthy participants. We define a voxel’s hemodynamic “delay” as the time a BOLD amplitude change occurs with respect to an amplitude change in a reference signal (also called a ‘probe regressor’). These are *relative* hemodynamic delays, representing the timing response of one brain region relative to another; this is therefore not directly capturing arterial transit times, though regional differences in transit times may affect these estimates. Rather than emphasizing local neural activity patterns, which may or may not relate to all brain regions, the probe regressor is commonly a physiological signal reflecting a more systemic fluctuation. The probe may be derived from an external recording related to cardiac or respiratory-related phenomena. Alternatively, a data-driven probe regressor can be used, typically the average BOLD signal in brain or grey matter tissue filtered to a certain frequency band, which is thought to mostly reflect these global systemic drivers (Aso et al., 2017; Liu, Li, et al., 2017; Tong et al., 2019). Previous work has also shown maps of hemodynamic delays to have good inter-session reproducibility and be more stable across different experimental designs when using a recursive tracking method for modelling (as implemented in this paper), versus a simple seed-based cross correlation approach (Aso et al., 2017).

Using this data-driven strategy, we assess the agreement of voxel-wise hemodynamic delay estimates derived from data that contained a breath-hold task and from resting-state data, to understand how similar these delay estimates are within an individual. To further interpret our assessment of the agreement, we applied amplitude thresholds (Chang et al., 2008; Fesharaki et al., 2021) and different levels of spatial smoothing of the input data, hypothesizing that signal-to-noise factors would modulate the absolute agreement of hemodynamic delay estimates. We also investigated which areas of the brain (if any) had the highest agreement, to help provide anatomical insight into what may be driving agreement or disagreement. Comparing these two paradigms may shed light on their different physiological drivers, or suggest they provide similar information under certain methodological constraints.

## 2. Methods

The data from this study unfortunately cannot be made openly available due to restrictions of the ethical approval that they were collected under. The visual instructions for the breathing tasks, displayed during scanning, were created with PsychoPy code (Stickland, Bright, Moia, & Zvolanek, 2022) and the hemodynamic timing estimates were produced with the *Rapidtide* Toolbox v2.0.9 (Frederick, 2016-2022). Specific details on how these coding repositories were used, as well as other primary analysis code for this project, have been organized into this GitHub repository: github.com/BrightLab-ANVIL/Gong_2022.

### 2.1 Data acquisition

This study was reviewed by Northwestern University’s Institutional Review Board. Written informed consent was obtained from all subjects. Nine healthy volunteers were included in the study (6 females, mean age = 26.2±4.1 years).

A 3T Siemens Prisma MRI system with a 64-channel head coil was used to collect neuroimaging data. The functional T2*-weighted acquisitions run during the hybrid breathing-task and resting-state protocol (Figure 1A) were gradient-echo planar sequences provided by the Center for Magnetic Resonance Research (CMRR, Minnesota) with the following parameters: TR/TE = 1200/34.4 ms, FA = 62°, Multi-Band (MB) acceleration factor = 4, 60 axial slices with an ascending interleaved order, 2 mm isotropic voxels, FOV = 208 × 208 mm^2^, Phase Encoding = AP, phase partial Fourier = 7/8, Bandwidth = 2290 Hz/Px. One single-band reference (SBRef) volume was acquired before the functional T2*-weighted acquisition (the same scan acquisition parameters without the MB acceleration factor) to facilitate functional realignment and masking. A whole brain T1-weighted EPI-navigated multi-echo MPRAGE scan was acquired, adapted from (Tisdall et al., 2016), with these parameters: 1 mm isotropic resolution, 176 sagittal slices, TR/TE1/TE2/TE3 = 2170/1.69/3.55/5.41 ms, TI = 1160 ms, FA = 7°, FOV = 256 × 256, Bandwidth = 650 Hz, acquisition time of 5 minutes 12 seconds, including 24 reacquisition TRs. The three echo images were combined using root-mean-square.

**Figure 1.**
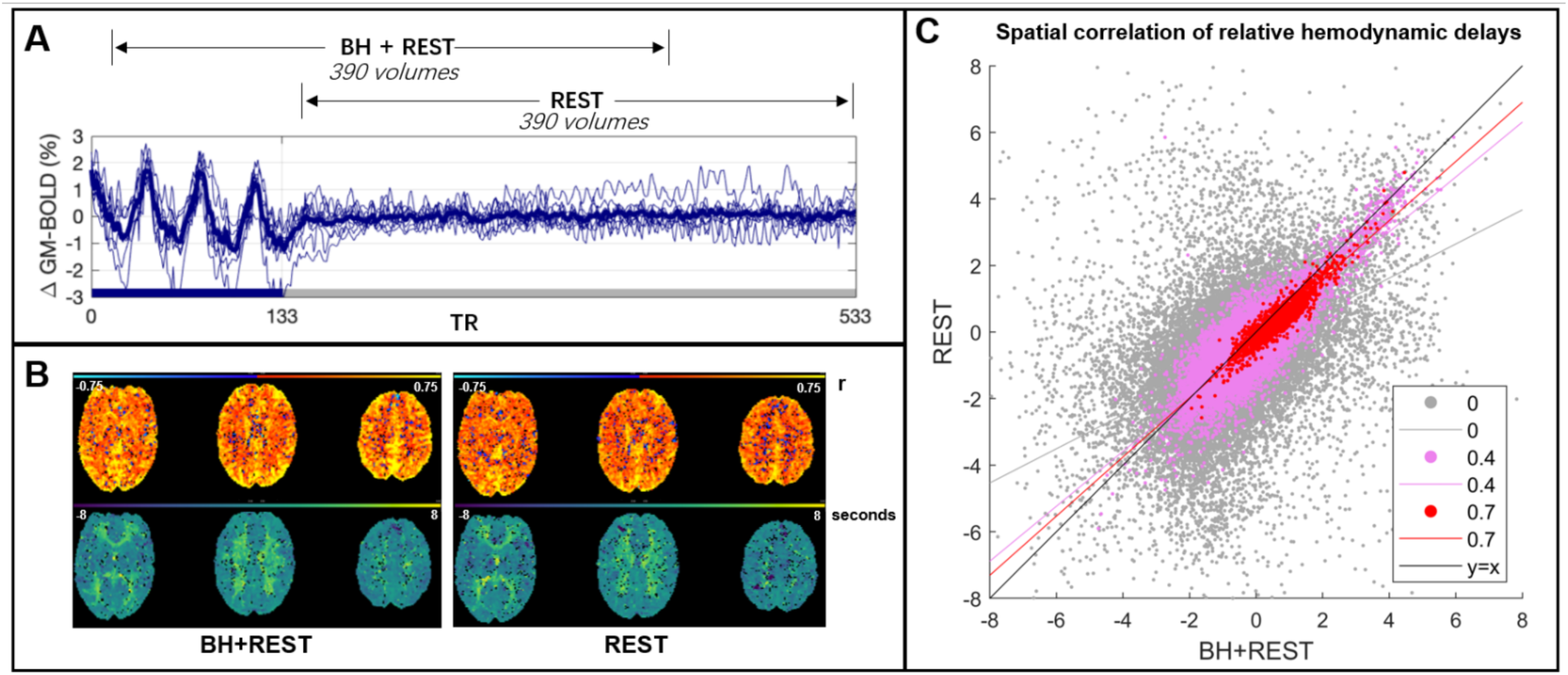
Panel A: The BOLD time-series averaged across gray matter (GM) for each subject (thinner blue lines) and the group average (thicker line). This time-series data was cut into two segments: one segment, BH+REST, includes a breathing task (solid blue bar on x-axis) followed by REST; the second segment is only REST. 390 volumes amount to 7 minutes 48 seconds (TR=1.2 seconds). Panel B: The output maps, from Rapidtide, for one example subject. The top row shows the correlation amplitude at each voxel (maximum correlation coefficient between each voxel time-series and probe regressor). The bottom row shows the relative delays at each voxel (time in seconds when the maximum correlation occurred). Panel C: Spatial correlation, in GM voxels, between hemodynamic delay times (panel B, bottom) from the BH+REST and REST data segments of an example subject. The correlation amplitude (panel B, top) is used to threshold the delays; the figure legend shows 3 examples of amplitude thresholding we applied. Each dot represents a voxel passing the threshold for both data segments, e.g., 0 = hemodynamic delay for a specific voxel must be in GM and have a correlation amplitude greater than 0 to be included. The lines are linear regression lines. For comparison, the y=x line is also shown.

Previous work with similar tasks (Bright et al., 2009; Lipp, Murphy, Caseras, & Wise, 2015) was used as guidance for the breath-hold (BH) task timings. Subjects practiced the BH tasks outside the scanner with the researcher (R.C.S) prior to scanning. During the scan, subjects performed three BH cycles (paced breathing and a 15-second end-expiration hold) followed by an 8-minute period of rest with visual fixation. Two equal-length data segments (Figure 1A) were analyzed: one comprised the breathing task followed by rest (BH+REST), whereas the other just included rest (REST). These neuroimaging data were used within previously published work (Stickland et al., 2021).

### 2.2 Data analysis

#### 2.2.1 Data preprocessing

A custom shell script grouped AFNI (Cox, 1996) and FSL (Jenkinson, Beckmann, Behrens, Woolrich, & Smith, 2012; Li, Morgan, Ashburner, Smith, & Rorden, 2016; Smith et al., 2004; Woolrich et al., 2009) commands. DICOMS were converted to NIFTI format with dcm2niix (Li et al., 2016). For motion correction of the fMRI dataset, AFNI’s 3dvolreg was performed with the SBRef as the reference volume; six motion parameters (three translations, three rotations) were extracted, demeaned, and saved. The first 10 volumes were discarded to allow the signal to achieve a steady state of magnetization and then brain extraction (Smith, 2002) was performed using FSL. The T1-weighted dataset was processed with the fsl*_*anat function, involving brain extraction, bias field correction, and tissue segmentation with FAST (Zhang, Brady, & Smith, 2001). A GM tissue mask was subsequently created by thresholding the partial volume estimate image to 0.5 and transforming it to the subject-specific fMRI space with FSL’s FLIRT command (Jenkinson, Bannister, Brady, & Smith, 2002; Jenkinson & Smith, 2001). The fMRI dataset was portioned into two segments (BH+REST and REST), each containing 390 volumes (Figure 1A). Finally, detrending and denoising were performed via linear regression using AFNI’s 3dTproject command, including the removal of Legendre polynomials up to the 4th degree to remove drifts in the signal and the six motion parameters. This single-subject preprocessed fMRI dataset, divided into two equal-length segments, was the input to the *Rapidtide* function, explained below.

The MNI structural atlas (Collins, Holmes, Peters, & Evans, 1995; Kötter et al., 2001), which parcellates the brain into 9 anatomical regions (caudate, cerebellum, frontal lobe, insula, occipital lobe, parietal lobe, putamen, temporal lobe, thalamus), was linearly transformed to fMRI subject space using FSL FLIRT, then masked to the subject’s GM. This transformation was achieved using nearest neighbor interpolation and 12 degrees of freedom and concatenating previously created registration matrices (the functional to structural space registration matrix and the structural to MNI registration matrix were previously created from running the fsl_anat function).

#### 2.2.2 Estimation of relative hemodynamic timings

The *Rapidtide* toolbox (v2.0.9) a suite of Python programs, was used to create maps of relative hemodynamic timing. *Rapidtide* implements an algorithm called *RIPTiDe* (Regressor Interpolation at Progressive Time Delays) that performs rapid time delay analysis in filtered fMRI data to find lagged correlations between each voxel’s time-series and a reference time-series, referred to as a “probe regressor” (Frederick, 2016-2022). We used a data-driven probe regressor, initially defined as the mean GM time-series. We chose to use the *Rapidtide* toolbox to estimate these relative hemodynamic timings because: (i) it is a citable open-source software package, (ii) it has been demonstrated to be successful when applied to resting-state data (Champagne et al., 2022; Erdogan, Tong, Hocke, Lindsey, & Frederick, 2016) and (iii) with a data-driven probe regressor, the resting-state and breath-hold data can be analyzed with an identical pipeline.

The *Rapidtide* commands are included in the GitHub repository listed earlier in this manuscript, and the input arguments used are summarized in Supplementary Table 1. These commands were run separately on the BH+REST and REST fMRI data segments for all subjects. Two *Rapidtide* output maps were extracted for further analysis (Figure 1B): the hemodynamic delay (temporal offset of the maximum correlation with the probe regressor, in seconds) and the correlation amplitude (maximum similarity function value, Pearson’s r). *Rapidtide* resamples the probe regressor in each refinement step and produces correlation estimates on a continuous scale within the search range given. We chose a search range of –15 to +15 s to comfortably cover expected physiological ranges in a healthy cohort alongside using a data-driven probe regressor with no large offset. In our further analyses of delay values, we did not include any values at the boundary of this search range, deeming these not a true optimization and likely not physiologically plausible.

#### 2.2.3 Hemodynamic delay comparison

The voxel-wise agreement between delay values from BH+REST and REST was calculated for all GM voxels using linear regression analysis, via the *polyfit* function in MATLAB (MATLAB and Statistics Toolbox Release 2021b, The MathWorks, Inc., Natick, Massachusetts, United States). This was repeated for subsets of GM voxels with increasing correlation amplitude thresholds applied (step=0.1, max 0.7): a voxel’s correlation amplitude value had to pass the threshold in both BH+REST and REST data segments of a given subject to have the delay values included.

Three metrics were summarized: the spatial correlation coefficient (representing model fit), the slope of the linear regression model (representing absolute agreement in values), and the percentage of voxels remaining after amplitude thresholding with respect to the entire GM. When the amplitude threshold is labeled as ‘None’ (e.g., x-axes in Figure 2), the percentage of voxels remaining is not at 100% because *Rapidtide* fails to produce a hemodynamic delay estimate for select GM voxels, resulting in the initial exclusion of these voxels from analysis. Figure 1C demonstrates the relationship between delay values at three different amplitude thresholds, for one example subject. All three metrics were then summarized across subjects using boxplots. The three metrics were also investigated within subregions of GM voxels. Using the MNI atlas described above, (Section 2.2.1), the same linear regression was performed for each GM atlas region (or parcel), then a median value across subjects was calculated for each metric, in order to easily visualize and compare the pattern of results between GM subregions.

**Figure 2.**
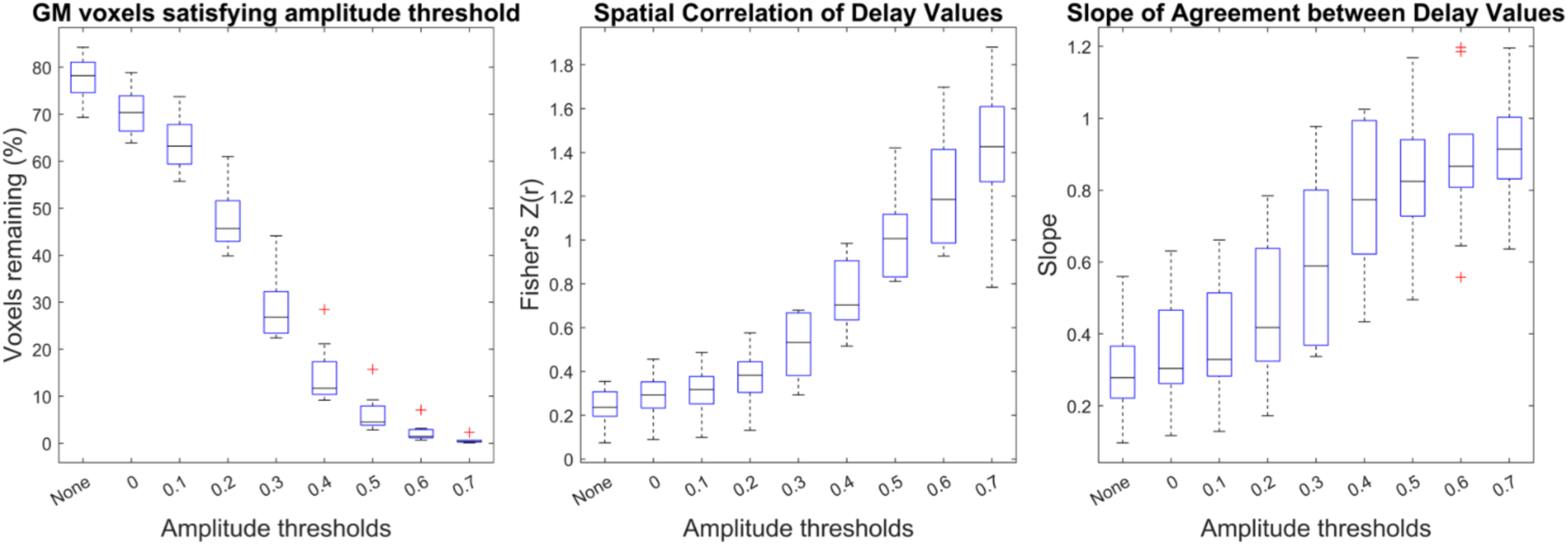
Delay agreement between BH+REST and REST data segments within GM voxels, for each amplitude threshold. The boxplots summarize across subjects. The left plot shows the percentage of GM voxels satisfying each amplitude threshold in both data segments, where ‘None’ refers to no amplitude thresholding applied, and the subsequent numbers (0 to 0.7) refer to the amplitude threshold the voxel must exceed in order to be included in the assessment of delay agreement. The middle and right plots summarize the delay agreement for each amplitude threshold, showing the spatial correlation coefficients (middle plot) and slopes (right plot).

#### 2.2.4 Impact of spatial smoothing

Prior research using a lagged-GLM approach to map regional hemodynamic timings showed that these results were more robustly achieved in breath-hold versus resting-state data (Bright & Murphy, 2017; Stickland et al., 2021), likely due to the reduced signal-to-noise of respiratory-evoked vascular responses in resting-state data. To evaluate the role of signal-to-noise (SNR) limitations in the current results, we enhanced the degree of spatial filtering applied to the data prior to hemodynamic delay estimation, from σ = 1mm (FWHM 2.35mm, *Rapidtide* recommended setting for smoothing) to σ = 2.13mm (5mm FWHM). The same three metrics were calculated and compared with the original less-smoothed results.

#### 2.2.5 Impact of overlapping time-series data

In the above analysis, the breathing task data segment and the resting-state data segment were extracted from the same functional scan acquisition for each of the subjects. In order to maximize and match the degrees of freedom between data segments, there is substantial overlap in the time-series information between the BH+REST and REST data segments (as indicated in Figure 1A). It is unclear whether this time-series overlap is critical to our observations of delay mapping. Therefore, in order to determine the effect of time-series overlap, an additional comparison of the BH+REST data segment against the REST data segment from another scan was conducted (Specifically, we isolated the resting state segment from the CDB+REST scan acquired in the same session, where “cued deep breathing” (CDB) is an alternative breathing task to BH, described in greater detail in Section 4.3.)

## 3. Results

### 3.1 Hemodynamic delay agreement between breathing-task and resting-state data

Figure 1B displays amplitude and delay maps for one example subject, and maps for all subjects are presented in Supplementary Figure 1. For both BH+REST and REST data segments, correlation amplitude values were generally positive and show tissue contrast between GM and white matter (WM). Voxels with negative correlations, or where *Rapidtide* was not able to output a value, appear predominantly in or adjacent to the cerebrospinal fluid (CSF) and sometimes in WM; this is more notable for the REST data segment. The REST data segments also display lower correlation amplitudes across the brain compared to BH+REST, as anticipated. Delay maps show good GM/WM contrast for both data segments, although some voxel clusters of extreme values are present (see Discussion).

Figure 2 shows the summary of the three metrics used to assess hemodynamic delay agreement at different correlation amplitude thresholds: percentage voxels remaining, spatial correlation, and linear slope. As the amplitude threshold increases, fewer voxels remain, and the spatial correlation and slope of agreement relating the BH+REST vs REST hemodynamic delays both increase in a non-linear manner (Figure 1C, Figure 2). In other words, the agreement between GM hemodynamic delay times in REST and BH+REST increases with stricter amplitude thresholding, as shown by the higher spatial correlation and the slope approaching a value of 1. Scatterplots comparing BH+REST and REST delay values for select amplitude thresholds are provided for all subjects (Supplementary Figure 2).

The parcellation analysis showed similar trends in each brain region examined (Figure 3). For each brain region, the percentage of voxels remaining (for each data segment separately and their union), the spatial correlation and the slope all changed similarly with increasing amplitude threshold.

**Figure 3.**
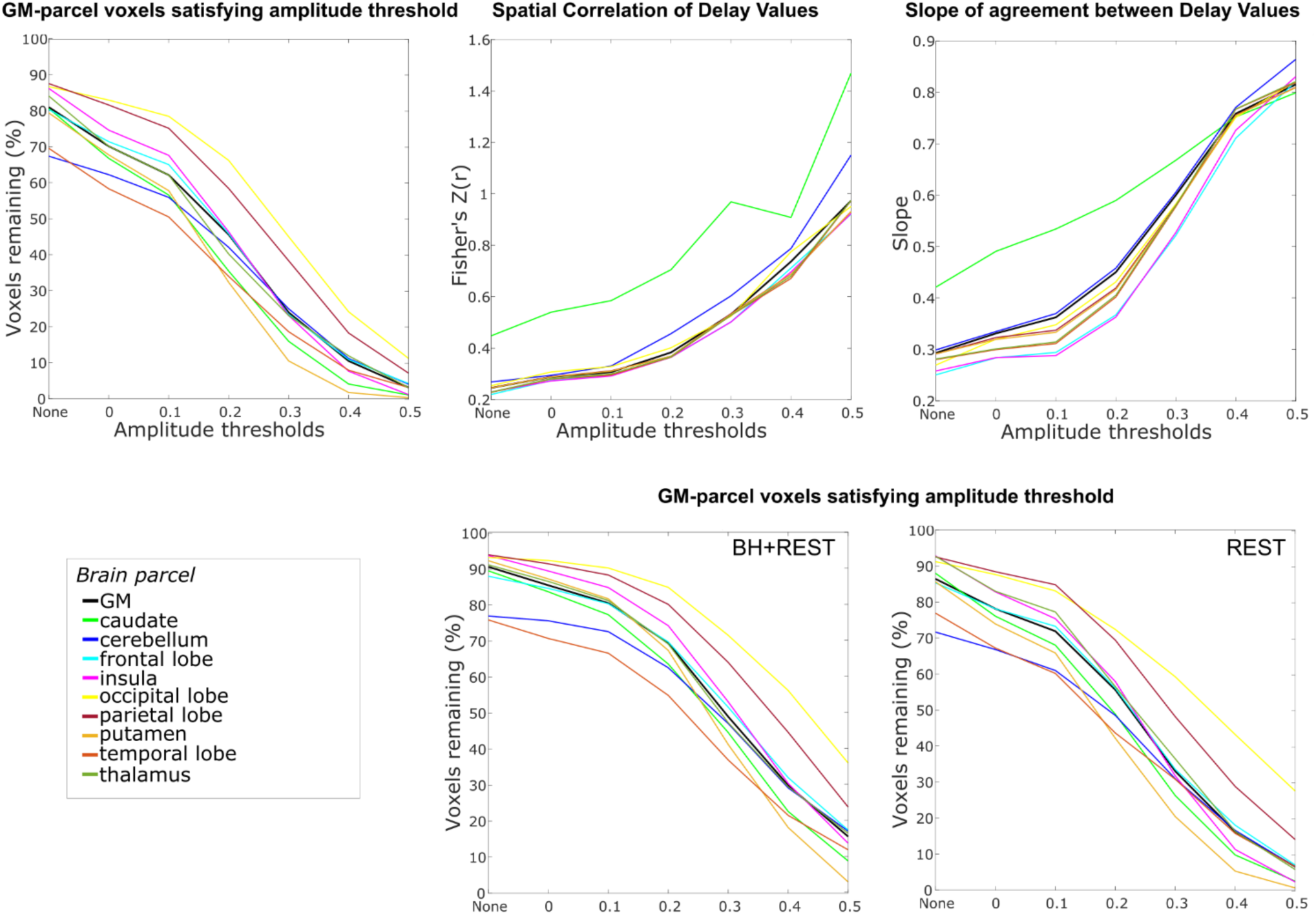
Delay agreement between BH+REST and REST data segments within GM parcel voxels, for each amplitude threshold. The three plots on the top row are displayed in the same format as Figure 2 but display the median value across subjects for each GM parcel (colored lines) alongside the median value for total GM (black line), for visualization simplicity. Additionally, the bottom row shows the percentage of voxels remaining for both data segments separately. The highest amplitude threshold that is summarized is capped at 0.5 because, for some GM parcels and some subjects, too few voxels pass the higher amplitude thresholds.

After visualizing the spatial location of voxels passing high amplitude thresholds (0.6, 0.7), we noted their proximity to the sagittal sinus and other regions that are expected to have large venous blood vessel contributions to the fMRI signal (Figure 4). Based on this observation, we hypothesized that higher amplitude thresholds may isolate voxels reflecting larger veins rather than tissue. The voxel-wise Mean Signal Intensity (MSI) is partially reflective of local venous weighting: venous blood induces large signal dephasing, resulting in lower MSI across the functional BOLD-weighted images. These voxels may drive agreement between BH+REST and REST delay values, without faithfully representing tissue physiology.

**Figure 4.**
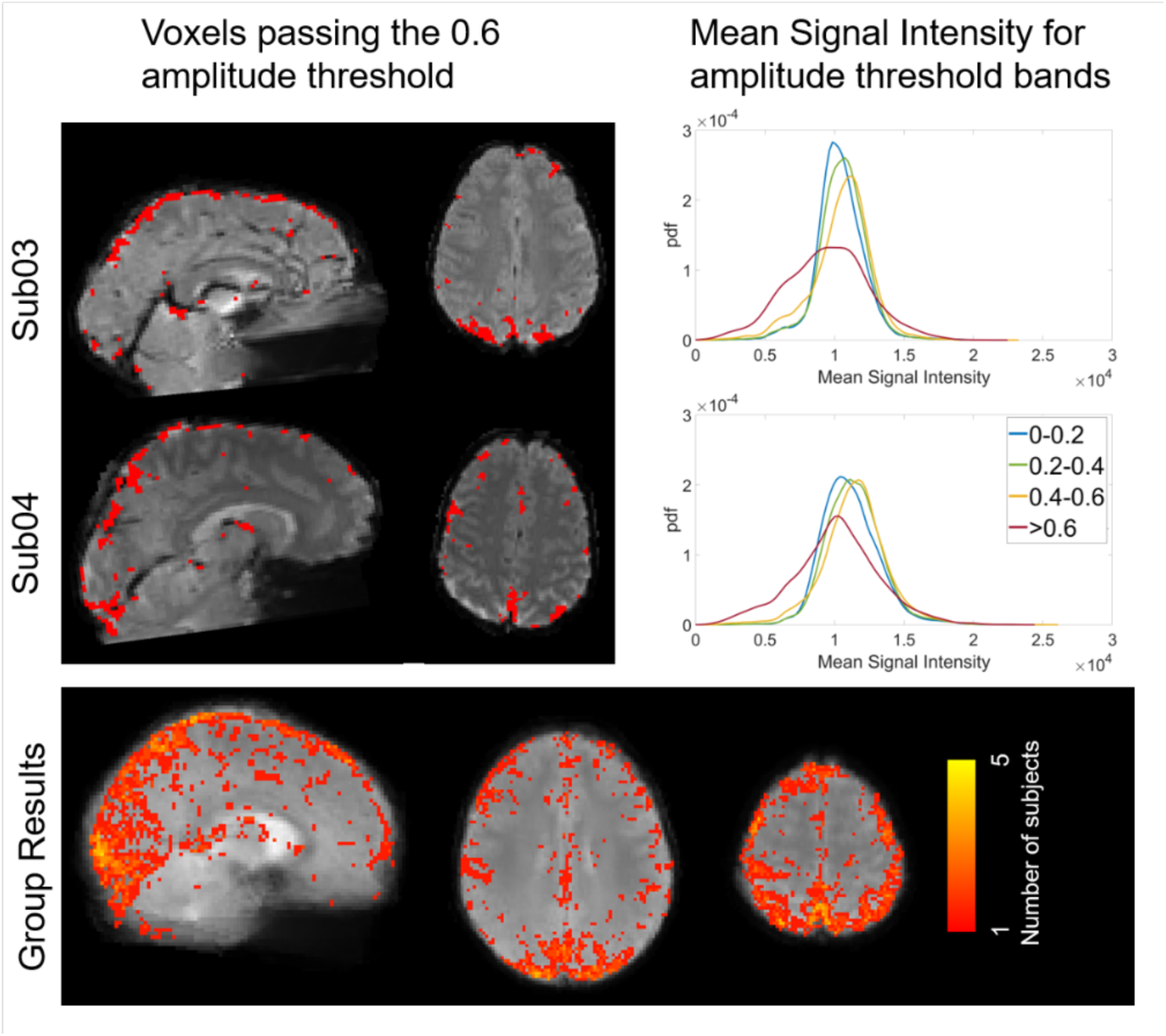
Voxels passing the 0.6 amplitude threshold in both BH+REST and REST data segments overlaid on mean signal intensity (MSI) maps, and histograms showing MSI distributions for different amplitude threshold bands, for two example subjects and group-level summary. Example subject maps are in subject space and the group result maps are in MNI space

To address this, we investigated the relationship between MSI and correlation amplitude threshold, hypothesizing the voxels passing the higher thresholds in both BH+REST and REST data segments would have lower MSI than the other thresholds, potentially due to the presence of larger veins. To visualize, MSI maps were overlayed with voxels passing the 0.6 amplitude threshold for both single subject and group data (Figure 4). Using the *ksdensity* function in MATLAB, the histograms of MSI values of voxels corresponding to each of the amplitude threshold bands 0-0.2, 0.2-0.4, 0.4-0.6, and larger or equal to 0.6 were extracted (Figure 4, with results from all subjects shown in Supplementary Figure 3). We observed that voxels with the highest correlation amplitudes in both data segments (r > 0.6) did generally reflect lower MSI values than other GM voxels, supporting our hypothesis that these voxels, at least in part, contain larger venous vessels.

To determine whether these voxels with very high correlation amplitudes drive the pattern of results presented in Figure 2, the original analysis was repeated for correlation amplitude bands rather than lower thresholds: 0 to 0.2, 0.2 to 0.4, and 0.4 to 0.6. However, with these amplitude threshold bands, only voxels within the threshold band for both data segments can be included, which ultimately resulted in only ∼25% of total GM voxels being compared in subsequent analyses (Supplementary Figure 4). This could have originated from the higher SNR of the BH+REST data segment compared with that of the REST task, such that global vasodilatory effects led to systematically higher correlation amplitude values. We therefore standardized correlation amplitudes in each data segment into percentile scores, and examined voxels that were in the same quartile for both data segments in an attempt to include more GM voxels in the subsequent analysis. Scatterplots showing these relationships for all subjects are provided in Supplementary Figure 5. Figure 5 shows the summary of the three metrics used to assess hemodynamic delay, revealing similar trends to the original results: the spatial correlation increases and the slope increases, towards 1, with each increasing amplitude percentile threshold band. Note that the percentages of voxels included in these threshold bands are still small after matching by percentile scores, indicating that many voxels show very different correlation amplitudes in the two data segments.

**Figure 5.**
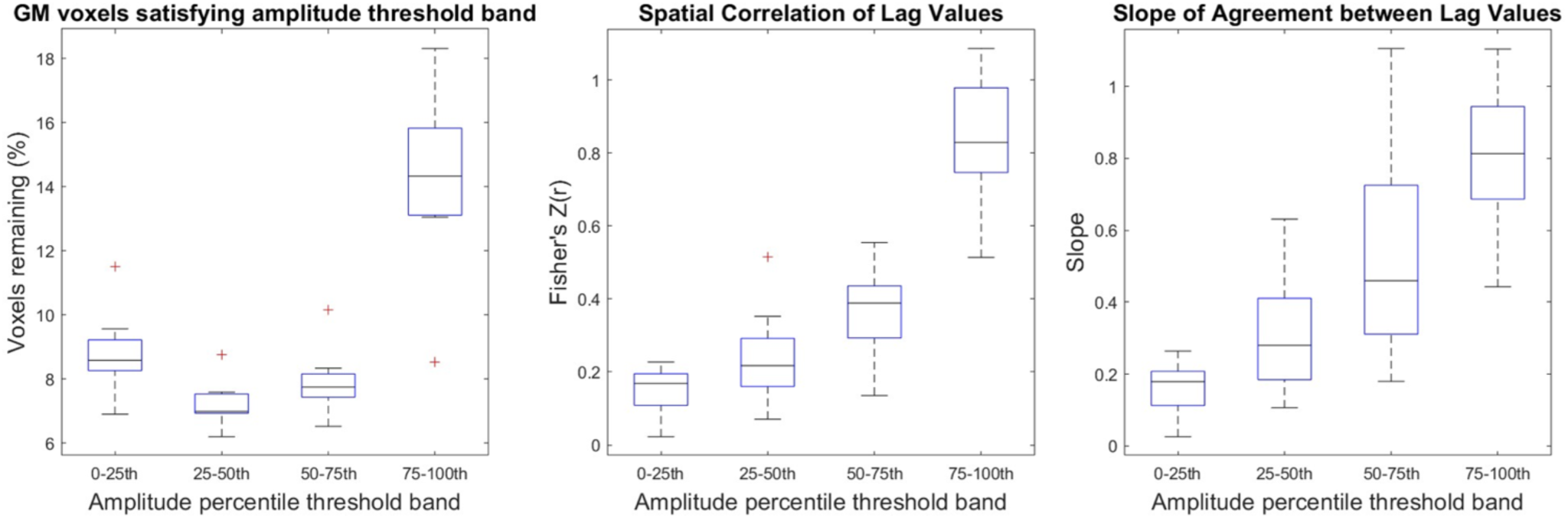
Delay agreement between BH+REST and REST data segments within GM voxels, for each amplitude percentile threshold band. The boxplots summarize across subjects. These results are displayed in the same format as Figure 2 (left: percentage of GM voxels satisfying each amplitude threshold band, middle: spatial correlation coefficients, right: slopes) but instead of filtering the voxels with a single lower amplitude threshold, amplitude percentile threshold bands are used (a voxel must be within the percentile threshold band in both data segments to be included).

### 3.2 Impact of Spatial Smoothing

Under additional spatial smoothing, we observe a widespread increase in correlation coefficient amplitudes (Figure 6) in both the BH+REST and REST data, which most likely reflects the improved signal-to-noise ratios achieved through this preprocessing step. Maps generated from the smoothed data from all subjects are presented in Supplementary Figure 6, and scatterplots comparing the resulting BH+REST and REST delay values for select amplitude thresholds and threshold bands are provided for all subjects (Supplementary Figures 7 and 8). Figure 7A demonstrates the effects of spatial smoothing on agreement between relative hemodynamic delay estimates, with the original results (median values from Figure 2) provided for comparison. Due to the increased correlation coefficient amplitudes in both data segments, the percentage of voxels remaining increased after additional smoothing. However, the spatial correlations of hemodynamic delays observed at each threshold are mostly unchanged. There is trend for better agreement (slope closer to 1) at lower amplitude thresholds in the additionally smoothed data (Figure 7A), however this is not readily interpretable given the inter-subject variability and the small sample size available. For the amplitude threshold band analysis after additional smoothing (Figure 7B, Supplementary Table 2), we see a different pattern in the voxel percentages in each threshold band. This corresponds to our observation that, with additional smoothing, more voxels pass higher amplitude thresholding. For every amplitude threshold band, we observe an increase in group median correlation amplitude and slope values after additional smoothing.

**Figure 6.**
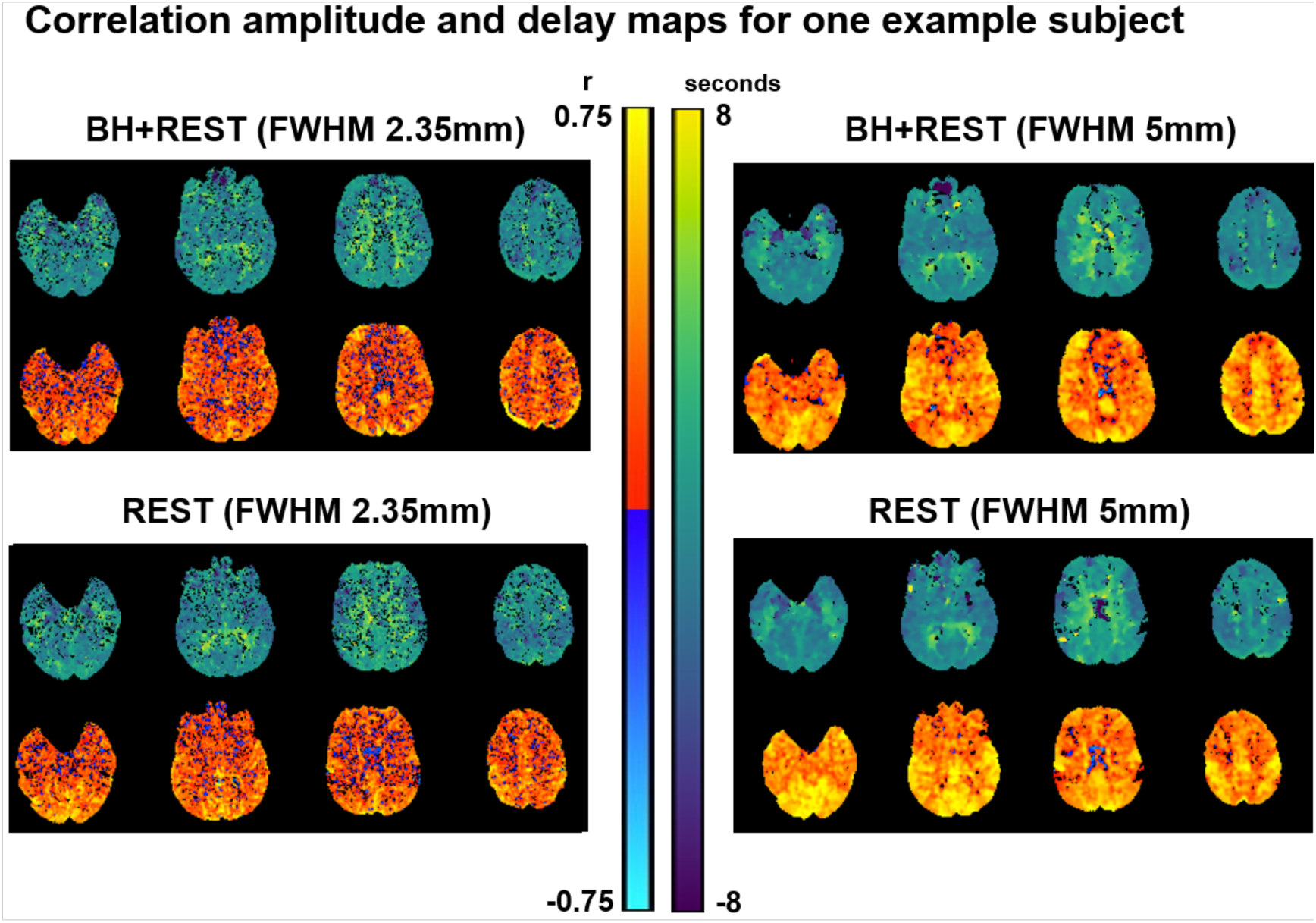
Parameter maps generated with Rapidtide, for one example subject (sub01), before and after changing the level of smoothing. The top row displays the relative delays at each voxel (time in seconds when the maximum correlation occurred). The bottom displays the correlation amplitude at each voxel (maximum correlation coefficient between each voxel time-series and probe regressor).

**Figure 7.**
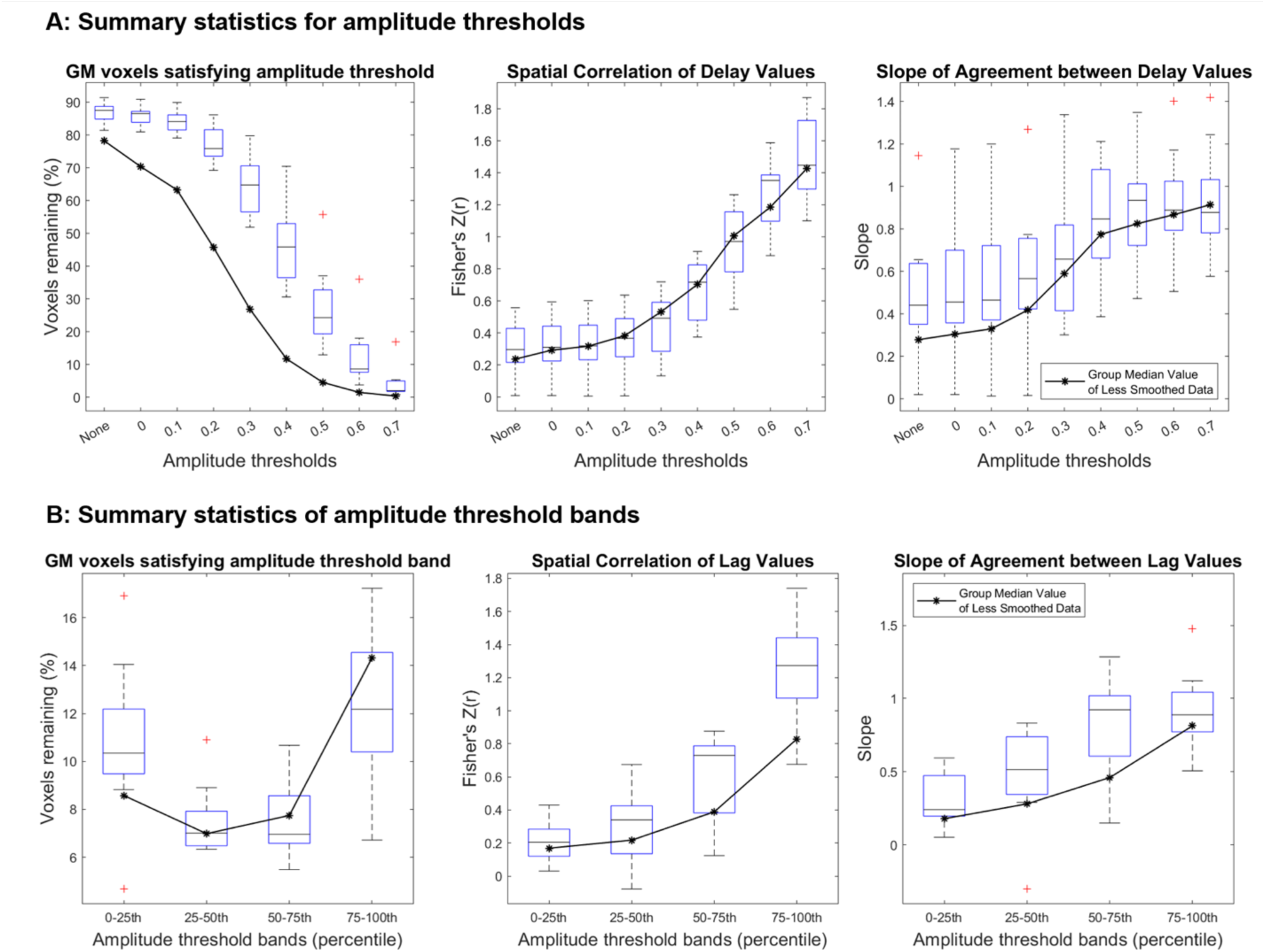
Panel A and B summarize amplitude threshold and amplitude percentile threshold band results, respectively, comparing the results with increased smoothing (FWHM 5mm) to the original results which had less smoothing (FWHM 2.35mm). The original results are shown as a black line for reference and duplicate the group median results from figures 2 and 5.

**Figure 8.**
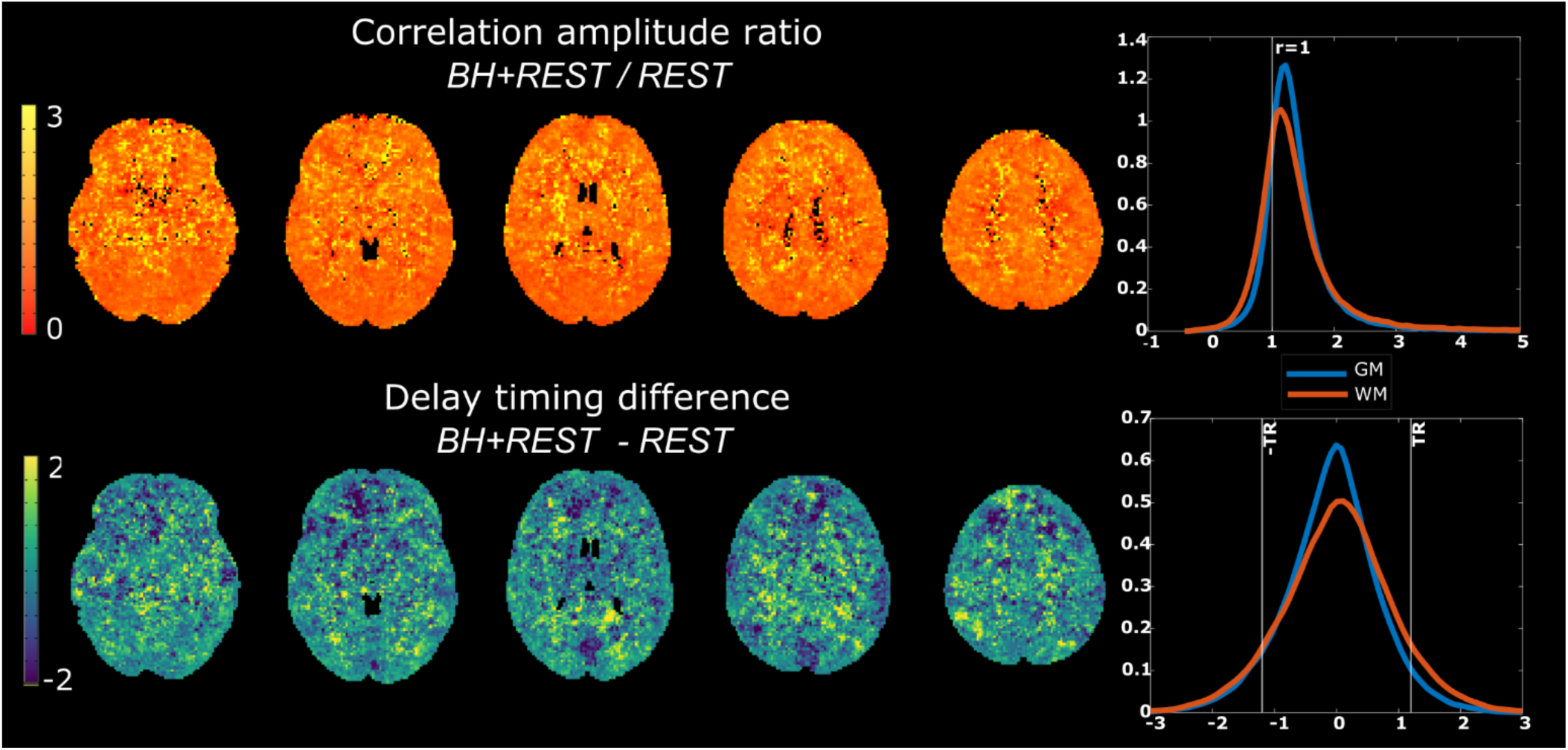
Top row: Ratio of the average correlation amplitude maps for BH+REST and REST data segments (Median: GM 1.27, WM 1.24); Bottom row: Average hemodynamic timing difference maps, BH+REST - REST (Median: GM -0.05, WM 0.04). In both rows, kernel density plots are provided to summarize gray matter (GM) and white matter (WM) values.

### 3.3 Impact of overlapping time-series data

The BH+REST and REST data segments are derived from the same scan and contain substantial overlapping time-series information. We examined the impact of this overlap on hemodynamic timing agreement, by comparing the BH+REST data segment with the REST segment from an entirely distinct functional acquisition (described in more detail in Section 4.3). The results are presented in Supplementary Figure 9. We observe the same effects as in our original analyses, suggesting that the agreement in hemodynamic timings is not entirely driven by the overlap in data segments.

## 4. Discussion

In BOLD-fMRI data containing a breath-hold task, we assume the globally driven vasodilatory effect dominates local hemodynamics, leading to a stronger relationship between a GM “probe” time-series and each voxel time-series. In comparison, in data containing only spontaneous resting-state fluctuations, the signal amplitude and timing characteristics at each voxel will be more region-specific, influenced more by neurovascular factors which are not globally driven. In this study, we modeled resting-state and breathing-task data segments with an identical analytical pipeline, to assess whether they produced equivalent estimates of relative hemodynamic timings across cortical brain regions. As we applied stricter correlation amplitude thresholds (correlation between the voxel time-series and the probe regressor), including a smaller percentage of the total GM voxels with each new threshold applied, the spatial agreement of hemodynamic delays between BH+REST and REST increased. Broadly, the same pattern of results was observed both when assessing all GM together and specific GM regions separately. We further observed that voxels passing the highest amplitude thresholds were more likely to be in closer proximity to venous structures; when removing these voxels from our analysis, we found similar trends to the original results, suggesting the delay agreement was not only driven by voxels proximal to large veins. Finally, applying additional smoothing to the input data boosted SNR and enhanced common variance shared across neighboring voxels; this produced widespread increases in correlation coefficient amplitudes, allowing more voxels to be included in the assessment of delay agreement. After additional smoothing, the pattern of agreement in delay values (as stricter thresholds were applied) did not change, yet some improvement in delay value agreement was observed at the group average level. The subtleties of these results, alongside their limitations and implications, are further discussed.

### 4.1 Estimating hemodynamic delays

Our implementation of the *Rapidtide* toolbox estimated a delay value for the majority of voxels in the brain, with the only inputs being the fMRI data and a GM tissue mask generated from a T1-weighted image. Therefore, this is a feasible and accessible way for researchers to map hemodynamic delays and can be implemented with the imaging data alone (i.e., without the need for additional equipment or recordings). The majority of the *Rapidtide* settings we applied are default or recommended by the toolbox developers with some input arguments adapted specifically for the nature of our dataset and questions, as described in the methods section and Supplementary Table 1. In Tong et al., 2019, the *Rapidtide* authors explain some of the features of using a global mean signal as the probe regressor: it is simple to implement, does not require an externally derived regressor, and resembles the signal fluctuations of the superior sagittal sinus (aligned with our observations in Figure 4). This will be true for the GM average signal as well, which we used to define and refine the probe regressor in this study. We chose to focus on GM to match the voxels where agreement of hemodynamic delay values was assessed, acknowledging that GM tissue has the most relevance to fMRI analyses that rely on neurovascular coupling. Furthermore, we did not want timing differences between GM and WM voxels to dominate our assessment of agreement between data segments.

In fMRI analyses, the global signal has traditionally been used within the context of denoising, with the reasoning that it captures global “noise” rather than local neural “signal”. However, its use is controversial considering the multitude of sources that can contribute to this signal (Liu, Nalci, & Falahpour, 2017). With the same logic that removing the global signal allows for the remaining variance to more closely resemble neural-driven mechanisms, specifically studying the global signal and its relationship to local signals can tell us more about non-neural hemodynamics. Tong et al (2019) discuss the challenges of using a data-driven mean signal regressor for both purposes; it is temporally “blurred” because it contains contributions from voxels with a range of delays and will also be influenced by local neural activity. To address these challenges *Rapidtide* implements a cross-correlation approach, often referred to as dynamic global signal regression in a denoising context (Erdogan et al., 2016). Within this cross-correlational approach, the *Rapidtide* algorithm goes through multiple analytical “passes”, a bootstrapping method to improve the probe regressor and make it more closely resemble non-neural processes (see the flow chart in Figure 5 of Tong et al., 2019). Further features, such as temporal filtering and iterative depseckling (using information from neighboring voxels to correct potentially bad delay estimates) are also applied. Please refer to Tong et al (2019) and the *Rapidtide* documentation (Frederick, 2016-2022) for more details and a more in-depth discussion of how voxel-wise hemodynamic delays are estimated and interpreted within this software package.

From visual inspection, the *Rapidtide* amplitude and delay maps broadly agree with previous CVR literature in healthy cohorts using resting-state or breathing-task designs, showing contrast between GM and WM tissue (Moia et al., 2021; Stickland et al., 2021; Tong et al., 2019), with negative CVR-related correlation amplitudes scattered more commonly across WM than GM but mostly clustered adjacent to CSF (Bright, Bianciardi, de Zwart, Murphy, & Duyn, 2014; Thomas et al., 2013). For some subjects, the delay maps contain clusters of extreme delay values (Supplementary Figures 1 and 6). Obvious explanations are unclear, but the *Rapidtide* despeckling procedure, and the smoothing we applied, could have accentuated these extreme clusters. They are present in both data segments but are more prominent in the REST data segment. Considering some of these regionally specific extreme delays, we also investigated the relationship between delay values at the level of GM cortical parcels. This allowed us to assess whether it was valid to summarize the patterns of agreement in delay at the level of the whole GM, or if different brain regions displayed notably different patterns of agreement across amplitude thresholds. However, we did not observe a meaningful difference in hemodynamic delay agreement across cortical regions.

### 4.2 Assessing the agreement of hemodynamic delays at different amplitude thresholds

For all subjects, we see positive correlations and slopes between BH+REST and REST data segments, at every amplitude threshold. The median slope across subjects, for every amplitude threshold, is below 1. This suggests a larger range of delay estimates in the data segment containing the breathing-task compared to the data segment containing resting fluctuations only. The results and relationships we observe across all GM voxels are consistent with those observed in the smaller GM subregions. It is unclear why the caudate stands out, with a slightly different pattern compared to other regions, though it still follows the same overall trend of increasing correlation amplitude and slope across increasing thresholding. To better visualize the agreement (or disagreement) between data segments, Figure 8 presents the ratio of the average (across subjects) correlation amplitude maps and average difference of the hemodynamic delay maps, using a visualization methodology similar to previous literature (Tong, Bergethon, & Frederick, 2011). It is clear that correlation amplitudes are systematically higher (ratio > 1) in the BH+REST data segment, as expected and visible in earlier figures. Interestingly, the hemodynamic delay differences are centered near zero, suggesting no systematic offset between the data segments, and the average distribution of this “error” in hemodynamic delay falls primarily within ±1TR for both GM and white matter (WM) voxels.

Further work is needed to concretely establish the reasons for (dis)agreement between hemodynamic delays measured in the breathing-task and resting-state data: are the relationships we observe driven by data quality, methodology, physiology, or a mixture of these factors? We attempted to explore these questions by assessing how the agreement changed when: (1) restricting the voxels included (via amplitude thresholds), (2) investigating specific voxels that may have a strong influence on the overall agreement and (3) changing the signal-to-noise ratio of the input data via increased spatial smoothing.

Firstly, if we examine the agreement of delay values with no amplitude threshold applied, therefore including most of the GM voxels within the linear model, we see very low correlations and slopes of 0.278 and 0.440 (median across subjects, less-smoothed and more-smoothed analysis, respectively). Moderate to strong correlations, with slopes closer to one, are only achieved when including a very small percentage of GM voxels, the voxels with the strongest positive correlation with the probe regressor. As a complementary representation of these results, we have also provided a summary of the root mean square error (RMSE) of the linear fits (Supplementary Figure 10). We show that RMSE decreases, meaning that “goodness of fit” increases, at higher amplitude thresholds, mirroring the correlation coefficient results. It is more difficult to interpret slopes when the correlation values are low, however the monotonic increase of correlation and slope, as amplitude threshold increases, is interesting to reflect on. With these results, we cannot conclude that the delay values estimated with resting-state and with breathing-task data have the same physiological meaning in every voxel; estimates may only converge when the weighting of neural, vascular and other physiological factors are the same. The improved delay agreement with higher amplitude thresholding may also reflect improved SNR, where higher correlation amplitudes make the measurement of hemodynamic delay more robust in the presence of other ongoing signal processes (“noise”).

Secondly, our results suggest that voxels with the strongest relationship with the probe regressor (in both data segments) are located close to the sagittal sinus and other regions that are expected to have large venous blood contributions. A similar observation was noted by a different group of researchers (Tong et al., 2011); they found, as have others, that the largest BOLD CVR-related amplitude changes are in large blood vessels, and that signal amplitude changes were most similar between a resting-state and a breath-hold dataset in these large vessels compared to in other regions. We went on to exclude these voxels in an attempt to assess hemodynamic delays relevant to local physiology, not just global vascular drainage. Without the inclusion of these high-amplitude voxels (within the upper quartile), we still observe a monotonic increase of spatial correlation and slope with each increasing amplitude percentile threshold band. This provides further evidence indicating that voxels with large venous vessels drive some of the agreement seen, but likely not all. Larger samples, and data acquisitions more tailored towards identifying venous weighting, would be needed to understand if this is meaningful.

Finally, increased smoothing of the input data increased the amplitude correlation coefficients of GM voxels on average (i.e., increased their correlation strength with the probe regressor). This is likely because of a reduced contribution from neurovascular fluctuations specific to each voxel, as well as other sources of local fMRI noise, leaving more shared features between neighboring voxels. This increased smoothing allowed more GM voxels to be included in the assessment of delay agreement for every amplitude threshold and manifested in similar or slightly improved hemodynamic delay agreement between the resting-state and breathing-task datasets. Therefore, this suggests that SNR factors do play a role in the agreement of delay estimates, and this is also consistent with the idea that higher agreement is observed in voxels with a stronger relationship to the global probe regressor. However, smoothing the data also amplified some unusual disagreement in hemodynamic delay observed in select subjects. For example, the scatterplots for Subject 3 in Supplementary Figures 2A and 7 demonstrate that there are select groups of voxels with near-zero hemodynamic delay in the BH+REST data but much more extreme relative delays in the REST data segment. (Note that this phenomenon appears limited to a small number of voxels, and it is counter to our observation, described above, that breathing-task data generally produces a wider range of delay values across all GM.) Such disagreement could suggest that the *Rapidtide* algorithm is problematically impacted by other ongoing signal fluctuations present in resting-state data that have distinct dynamics to low-frequency systemic vasodilatory effects and are coordinated across multiple contiguous voxels. In contrast, the higher SNR and greater specificity to CO_2_-related vasodilatory effects that are achieved by having a breathing task may prevent this fitting challenge, irrespective of spatial smoothing.

#### 4.3 Alternative breathing tasks

As an alternative to the BH task which induces hypercapnia, the neuroimaging dataset also included a cued deep breathing (CDB) task in the same participants, which induces hypocapnia via hyperventilation, described in (Stickland et al., 2021). Two repetitions of fast deep breaths were cued, and an 8-minute rest period followed this CDB task. As with the BH+REST dataset, two equal-length data segments were extracted. We replicated the main analysis with this alternative breathing task (the agreement of hemodynamic timings between CDB+REST and REST, across amplitude thresholds), shown in Supplementary Figures 2B and 11. Except for one subject being removed from group summaries as an outlier, we note the same trends and key findings reported in the main manuscript with the BH+REST dataset. Although our observations may be fairly robust to the breathing task used, we note that this may not be true in patients with vascular pathology; altered baseline vascular tone can differentially impact hypercapnic and hypocapnic cerebrovascular reactivity, including both the magnitude and delay of the vascular response (Bright, Donahue, Duyn, Jezzard, & Bulte, 2011; Zhao et al., 2009).

#### 4.4 Alternative reference signals

The complexity of the vascular tree, and the heterogeneous venous-weighting of BOLD fMRI contrast, makes it difficult to disentangle the fundamental biophysical meaning of the hemodynamic delays that we are measuring in our analysis. The reference time-series (e.g., probe regressor) used in our analysis is recursively derived from the GM mean time-series, which will reflect a mixed sampling of vascular sources. In the existing literature using similar delay mapping methodology, other groups have used a more purely venous reference time-series, derived from the superior sagittal sinus (SSS) (Aso et al., 2017; Tong et al., 2016; van Niftrik et al., 2016). Imitating their methods, we manually delineated an ROI in the SSS of each subject’s T1-weighted anatomical scan, registered this ROI mask to their functional data, and used this mask to define a probe regressor in *Rapidtide*. The correlation amplitude maps showed strong correlations and high agreement between GM and SSS results when compared within a GM ROI, particularly for the BH+REST segment (Supplementary Table 3). For delay map agreement, there was more subject variability, but on average these correlations are still moderate to strong, and again stronger for the BH+REST data segment. To bring context to strength of these correlations – no amplitude thresholding was applied within the GM ROI for the results presented in Supplementary Table 3, therefore this suggests greater similarity for maps produced with different reference signals (but the same data segment) compared to maps produced with the same reference signal (but different data segments), as shown in manuscript Figure 2 for GM with no amplitude threshold applied. We then replicated the original analyses described above, reproducing Figure 2 using the new SSS reference time-series results (Supplementary Figure 12). The trends we observe in this modified analysis (decrease in voxel percentage remaining, increase in Fisher’s Z spatial correlation, and increase in slope as the amplitude threshold is increased) are almost identical to those in the original results using a GM region to define the reference timeseries. Therefore, this change in reference time-series does not influence our main conclusions regarding hemodynamic delay timings in BH+REST versus REST data segments.

#### 4.5 Alternative approaches to estimating hemodynamic delays

There are alternative approaches to estimating hemodynamic delays with BOLD-fMRI data, each with their pros and cons relating to complexity of acquisition and analysis. Using probe regressors external to the brain such as end-tidal CO_2_ or near-infrared spectroscopy (Pinto et al., 2016; Tong et al., 2019), instead of a data-driven regressor, results in the regression coefficients having a more specific physiological interpretation and being easier to compare across studies. This is a more important priority when scaling BOLD amplitude changes, rather than timing (Zvolanek et al., 2022); timing estimates are often normalized to a common spatial reference (i.e., median timing across GM voxels, as done in this paper) even when an external probe regressor is used. Furthermore, we previously found that when using an external end-tidal CO_2_ regressor within a lagged general linear model framework it can be challenging to get trustworthy estimates of delay in resting-state data alone (Bright, Tench, & Murphy, 2017; Stickland et al., 2021). Other modeling approaches exist to extract timing information, for example, Fourier based analyses (Pinto et al., 2016), but without a signal model for the resting-state data, this cannot be applied in the same way as with a breathing task or gas delivery design with a specific task frequency. Compared to our task paradigm, previous papers using breathing tasks will typically employ more repeated breath-hold cycles to improve the modeling of vascular reactivity mechanisms, and it is also recommended to use multi-echo BOLD fMRI data with spatial ICA-denoising to increase BOLD signal-to-noise ratio and more effectively remove motion artifacts (Cohen & Wang, 2019; Moia et al., 2021). In our data, we regressed out the six motion parameters prior to running *Rapidtide*, however we cannot conclude that motion is fully accounted for. These factors should be considered to improve future comparisons of BOLD timing estimates from resting-state and breathing-task designs.

#### 4.6 Limitations and further considerations

The two data segments compared in this manuscript, BH+REST and REST, overlapped in time (Figure 1A). It could be argued that this would lead to artificial agreement when trying to compare ‘breathing-task’ data to ‘resting-state’ data; though this may be the case (despite contrary evidence presented in Section 3.3), our results show that good agreement of hemodynamic timings is only achieved under relatively strict analytical constraints, even with this overlap. Looking at Supplementary Figures 1 and 6, there are many clusters of voxels that show divergent hemodynamic delay estimates in BH+REST compared to REST (for example see subjects 07, 09, and 10), and much of the analysis we performed attempted to understand this disagreement. The advantages of using overlapping time segments include fewer confounding factors due to the participant being in a different physiological state. Furthermore, using two different sections of the same scan means sources of error driven by volume registration are minimized, making the voxel-wise comparisons more valid. This allowed us to more confidently interpret any differences in hemodynamic timing estimates as driven by replacing a resting-state portion of the dataset with a hypercapnic breathing task portion.

With our data and analysis methodology, we cannot specifically quantify arterial arrival times or blood flow changes in response to a known change in arterial blood gas tensions. However, when accounting for voxel-wise delays in fMRI analyses, and bringing clinical insight to regional cerebrovascular dysfunction, it is likely that relative hemodynamic timings are sufficient. We also anticipate that concurrent autoregulatory phenomena are present in both data segments, and that blood pressure fluctuations may be particularly enhanced in the BH+REST data segment and time-locked to the breathing task. Future work would benefit from the addition of beat-to-beat blood pressure monitoring to better characterize these changes and facilitate the differentiation of autoregulatory processes from neurovascular coupling or cerebrovascular reactivity effects. Furthermore, the short breath-hold tasks used in this study are known to drive mild hypoxia in addition to mild hypercapnia, and hypoxia may have distinct influence on the cerebrovasculature. The two effects are profoundly correlated in breath-holding fMRI data (Bright & Murphy, 2013), and difficult to tease apart in correlative time-series analysis. Thus, in our BH+REST segment, there are several concurrent effects causing BOLD signal changes: 1) vasodilation caused by increased PaCO_2_, 2) vasodilation caused by reduced PaO_2_, and 3) reduced PaO_2_ itself, all caused by the apnea. The two vasodilatory responses will both incur a BOLD signal increase throughout most of cortical GM, whereas reductions in PaO_2_ may lead to BOLD signal decreases. We note, however, that the vasodilatory effect of mild hypoxia is anticipated to be small (Brugniaux, Hodges, Hanly, & Poulin, 2007). Although we do not differentiate the contributions of these three mechanisms to our observations, we can make several inferences based on our data and the literature. First, Tancredi and Hoge (Tancredi & Hoge, 2013) reported that breath-hold BOLD fMRI signal changes did not substantially differ from signal changes associated with controlled gas inhalation challenges, where hypoxia could be minimized. Second, we observed similar results when considering BH+REST and CDB+REST data, and the CDB task has potential hyperoxic, rather than hypoxic, effects. Still, to determine the precise contributions of concurrent physiological fluctuations on the hemodynamic delay structure throughout the brain, future work would benefit from more controlled physiological stimuli to independently modulate PaO_2_ and PaCO_2_, or other time-locked physiologic effects.

The band-pass filter applied by the *Rapidtide* algorithm (0.0009 - 0.15Hz) should minimize the influence of the cardiac rhythm on our analysis, however aliased cardiac signals and other cardiac-mediated effects could still drive patterns in the lag structure particularly for participants with lower heart rates. Therefore, it may be worthwhile to specifically measure and model this alternative source of signal variation (to respiratory-mediated effects) in future studies. Furthermore, because the resting-state only data will have lower signal-to-noise compared to the data including the breathing task, these data may require additional noise removal to achieve more robust delay mapping. However, as the main comparison in this paper was to compare BH+REST with REST, we were hesitant to apply differing denoising strategies beyond the ones already implemented, particularly as a previous (single-echo) BOLD-fMRI study showed that delay maps lose both information and reproducibility when traditional ICA-denoising strategies are applied (Aso et al., 2017). Nevertheless, we do acknowledge that the absence of some of these denoising strategies in our resting-state data could have affected the level of delay agreement we observed. Future studies implementing multi-echo fMRI acquisition and denoising strategies could potentially help to mitigate this concern (Moia et al., 2021), while also adding insight into the biophysical mechanisms driving signal changes (Aso et al., 2019).

Finally, this study has a small sample size, and future work should investigate these questions further with larger and more diverse samples to determine generalizability. Additional data in patient cohorts with atypical regional hemodynamics would also help confirm the interpretation of our results. We provide many supplementary figures to complement the main results, in order to transparently communicate the full data variance in this healthy cohort.

## 5. Conclusions

We modeled BOLD fMRI data to estimate relative hemodynamic delay values across the brain, focusing on GM tissue. We investigated how these delay estimates change in the presence of a breathing-task versus in resting-state data. Hemodynamic delay information is a clinically meaningful marker of cerebrovascular function, and although breathing-task data may improve the robustness of delay mapping, resting-state data would not require the same degree of participant compliance and is therefore a very desirable approach. We found strong similarities between estimates from resting-state data and breathing-task data only when strict amplitude thresholds are applied, comparing a small percentage of GM voxels. A stronger agreement between delays estimated in these two different data types was seen in voxels that more closely resembled the global GM average, and localized (at least in part) to large venous structures. Caution must be taken when using voxel-wise delay estimates from resting-state and breathing-task data interchangeably throughout cortical GM, and further work is needed to more directly identify the local and global signal contributions that lead to their disagreement.

## Acknowledgments

This work was supported by the Center for Translational Imaging at Northwestern University. The authors express special thanks to Dr. Blaise Frederick for his assistance implementing the *Rapidtide* software package.

## CrediT author contribution statement

**Jingxuan Gong:** Conceptualization, Methodology, Software, Formal Analysis, Data Curation, Writing (OD), Writing (RE), Visualization, Funding Acquisition. **Rachael Stickland:** Conceptualization, Methodology, Software, Formal Analysis, Investigation, Data Curation, Writing (OD), Writing (RE). **Molly Bright:** Conceptualization, Methodology, Resources, Writing (OD), Writing (RE), Supervision, Project Administration, Funding Acquisition.

**Abbreviations**
*BH*: breath hold
*BOLD*: blood oxygenation level dependent
*CDB*: cued deep breathing
*CSF*: cerebrospinal fluid
*CVR*: cerebrovascular reactivity
*FC*: functional connectivity
*fMRI*: functional magnetic resonance imaging
*GM*: gray matter
*HRF*: hemodynamic response function
*MSI*: mean signal intensity
*WM*: white matter

