## Supplemental Material for "Hemodynamic timing in resting-state and breathing-task BOLD fMRI"

### Supplementary Material

| Rapidtide setting | Choice and rationale |
|---|---|
| delaymapping | A macro that sets multiple settings appropriate for a delay mapping analysis |
| passess | Set to 3 - a default set in the delay mapping macro. |
| despecklepasses | Set to 4 - a default set in the delay mapping macro. |
| refineoffset | Set to TRUE - a default set in the delay mapping macro. |
| pickleft | Set to TRUE - a default set in the delay mapping macro. |
| doglmfilt | Set to FALSE - a default set in the delay mapping macro. |
| searchrange | Set to –15 to 15. Physiologically plausible range of hemodynamic delays based on a healthy cohort and using a data-driven reference signal (probe regressor). |
| filterband | Set to lfo (0.0009-0.15Hz) - a default set in rapidtide. |
| datatstep | Set to 1.2, the TR of the input data. |
| detrendorder | Set to 0. Detrending was done prior to rapidtide. |
| oversampfac | Set to 4, instead of automatic factor (default) to ensure consistency across runs. |
| spatialfilt | Set to –1, recommended setting when performing smoothing in rapidtide (-1 refers to a sigma of half the mean voxel dimension. For our data, sigma was 1mm, which is 2.35mm FWHM. We re-ran with a sigma of 2.13mm, which is 5mm FWHM). |
| globalmeaninclude | Input was the GM tissue mask to exclude non-GM tissue in the initial calculations of the probe regressor, and to match the voxels where agreement of hemodynamic delay values was assessed. |
| refineinclude | Input was the GM tissue mask to exclude non-GM tissue in the refinement steps of the probe regressor, and to match the voxels where agreement of hemodynamic delay values was assessed. |

***Supplementary Table 1.*** *A summary of the key input arguments used for the Rapidtide algorithm.*

| FWHM (mm) | Data Segment | 25 | 50 | 75 |
|---|---|---|---|---|
| 2.35 | BH+REST | 0.2495 ± 0.0274 | 0.3641 ± 0.0404 | 0.4903 ± 0.0504 |
| | REST | 0.2008 ± 0.0231 | 0.2997 ± 0.0312 | 0.4137 ± 0.0425 |
| 5 | BH+REST | 0.4537 ± 0.0612 | 0.5755 ± 0.0576 | 0.6739 ± 0.0513 |
| | REST | 0.3262 ± 0.0595 | 0.4474 ± 0.0649 | 0.5682 ± 0.0663 |

***Supplementary Table 2.*** *Correlation amplitude value within GM tissue at 25th, 50th and 75th percentile, summarized across subjects (mean ± standard deviation). This is shown for each data segment (for each level of smoothing).*

| Agreement of Correlation Amplitude Maps (GM vs SSS) | | | | Agreement of Hemodynamic Delay Maps (GM vs SSS) | | | |
|---|---|---|---|---|---|---|---|
| BH+REST | | REST | | BH+REST | | REST | |
| Spatial Correlation (r) | Slope | Spatial Correlation (r) | Slope | Spatial Correlation (r) | Slope | Spatial Correlation (r) | Slope |
| 0.81 | 0.83 | 0.74 | 0.75 | 0.59 | 0.50 | 0.54 | 0.51 |
| 0.96 | 0.98 | 0.88 | 0.89 | 0.86 | 0.84 | 0.68 | 0.69 |
| 0.76 | 0.88 | 0.47 | 0.52 | 0.36 | 0.26 | 0.29 | 0.21 |
| 0.94 | 0.94 | 0.85 | 0.85 | 0.81 | 0.78 | 0.69 | 0.66 |
| 0.81 | 1.00 | -0.14 | -0.14 | 0.67 | 0.57 | 0.03 | 0.01 |
| 0.90 | 0.92 | 0.86 | 0.92 | 0.67 | 0.70 | 0.54 | 0.51 |
| 0.98 | 1.00 | 0.67 | 0.61 | 0.85 | 0.83 | 0.51 | 0.44 |
| 0.94 | 0.99 | 0.87 | 0.95 | 0.66 | 0.75 | 0.75 | 0.73 |
| 0.94 | 1.00 | 0.70 | 0.83 | 0.80 | 0.73 | 0.57 | 0.90 |
| **0.92±0.42** | **0.95±0.06** | **0.72±0.46** | **0.69±0.34** | **0.72±0.29** | **0.66±0.19** | **0.54±0.28** | **0.52±0.27** |

***Supplementary Table 3.*** *Agreement between the correlation amplitude maps (or delay maps) derived from the same data segment (BH+REST or REST) using either a gray matter (GM) or superior sagittal sinus (SSS) region of interest to define the probe regressor in the Rapidtide analysis. For all GM voxels, the spatial correlation (Pearson's r) and slope of a linear regression are reported for each subject and for the group average. Note, the correlation coefficients were transformed using the Fisher r-to-z transform prior to averaging and then reconverted.*

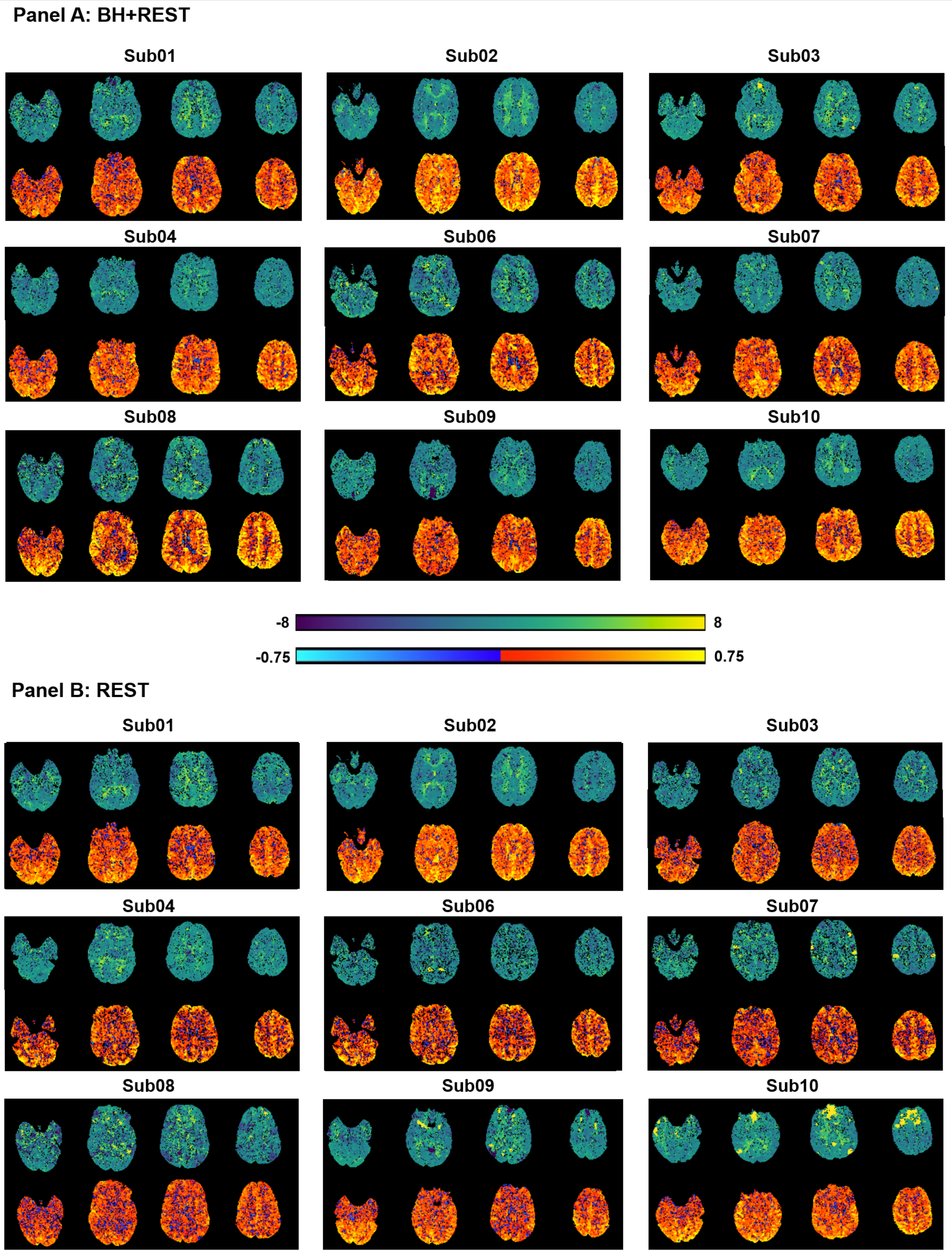


***Supplementary Figure 1.*** *Parameter maps generated with Rapidtide, for all subjects, for data segment BH+REST (panel A) and data segment REST (panel B). The top row for each subject and the top color bar displays the relative delay at each voxel (time in seconds when the maximum correlation occurred). The bottom row for each subject and the bottom color bar displays the correlation amplitude at each voxel (maximum correlation coefficient between each voxel time-series and probe regressor).*

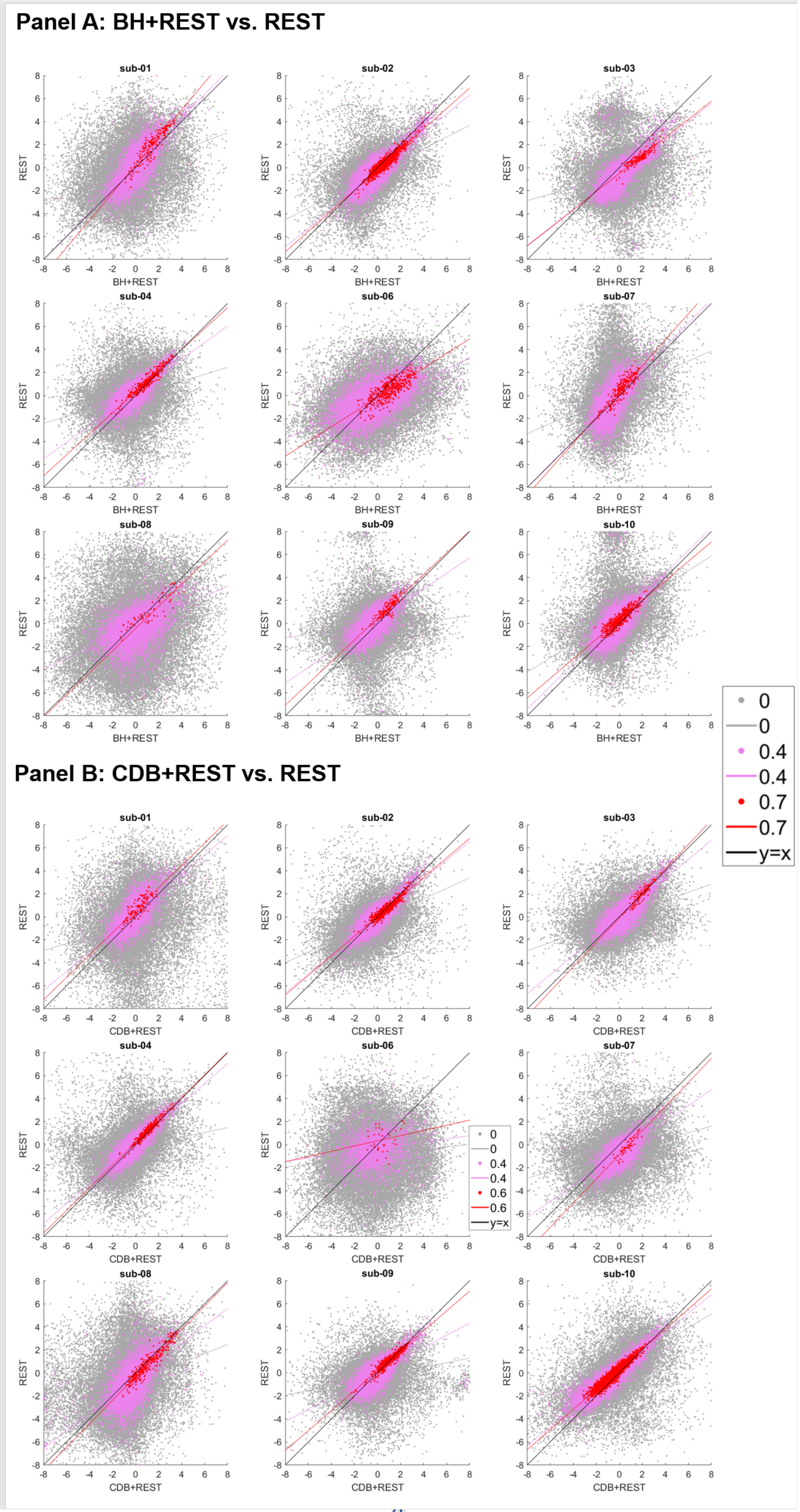

Panel A: BH+REST vs. REST
sub-01
sub-02
sub-03
sub-04
sub-06
sub-07
sub-08
sub-09
sub-10
REST
BH+REST
0
0
0.4
0.4
0.7
0.7
y=x
Panel B: CDB+REST vs. REST
sub-01
sub-02
sub-03
sub-04
sub-06
sub-07
sub-08
sub-09
sub-10
REST
CDB+REST
0
0
0.4
0.4
0.6
0.6
y=x

*(with previous page)*

***Supplementary Figure 2.*** *Spatial correlation, in GM voxels, between hemodynamic delay times from the TASK+REST and REST data segments, for all subjects. Panel A shows results from the breath-hold (BH) task, discussed in the main manuscript, and panel B shows the cued deep breathing (CDB) task. The CDB is different from the vasodilatory BH challenge and provides a vasoconstrictive stimulus. The correlation amplitude is used to threshold the delays; the figure legend shows 3 examples of amplitude thresholding we applied. Each dot represents a voxel passing the threshold for both time segments, e.g., 0 = hemodynamic delay for a specific voxel must be in GM and have a correlation amplitude greater than 0 to be included. The lines are linear regression lines. For comparison, the y=x line is also shown. For sub-06, panel B, there were too few voxels remaining after the 0.7 amplitude threshold therefore the highest amplitude threshold was set to 0.6.*

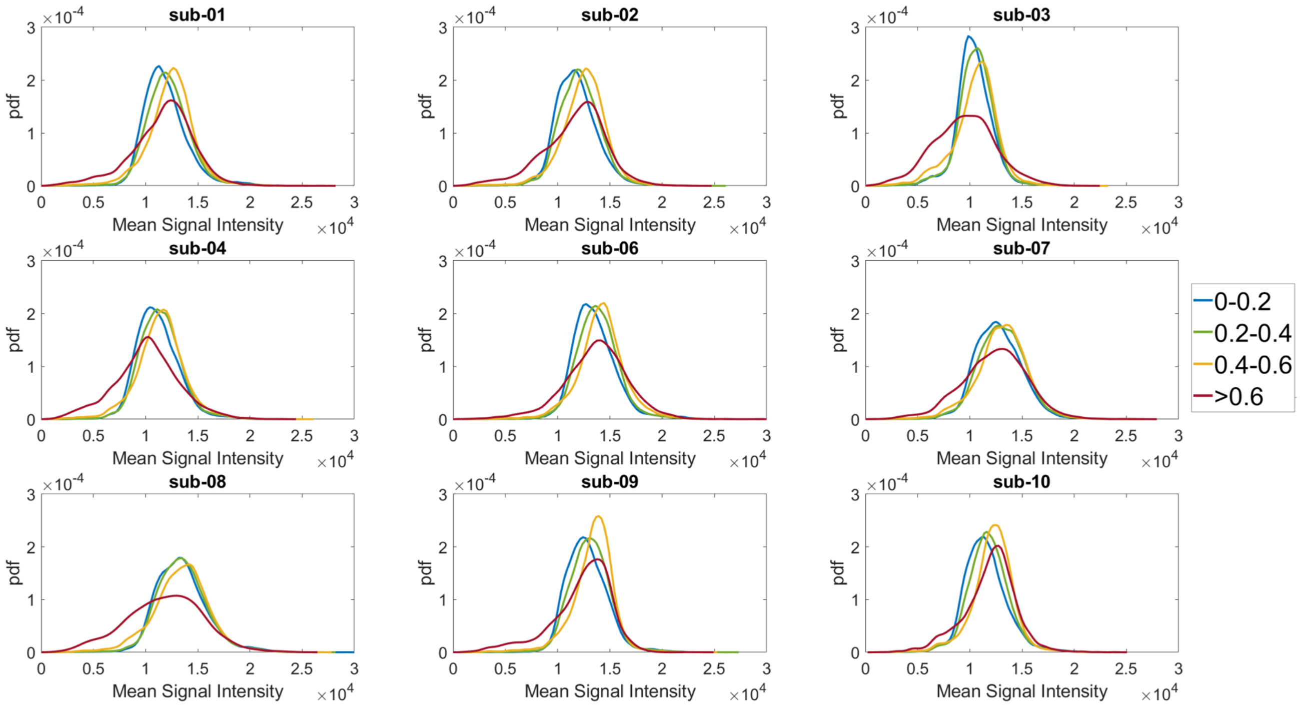


***Supplementary Figure 3.*** *Histograms, for each subject, showing the mean signal intensity (MSI) distributions for different amplitude threshold bands.*

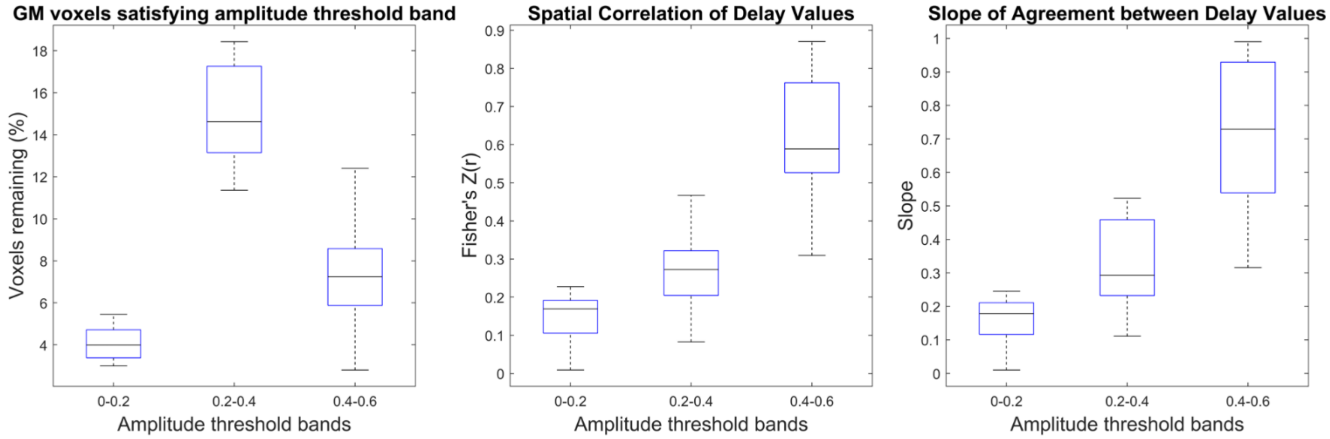


***Supplementary Figure 4.*** *Delay agreement between BH+REST and REST data segments within GM voxels, for each amplitude threshold band. The boxplots summarize across subjects. These results are displayed in the same format as Figure 2 (left: percentage of GM voxels satisfying each amplitude threshold, middle: spatial correlation coefficients, right: slopes) but instead of filtering the voxels with a single lower amplitude threshold, amplitude threshold bands are used (a voxel must be within the threshold band in both data segments to be included).*

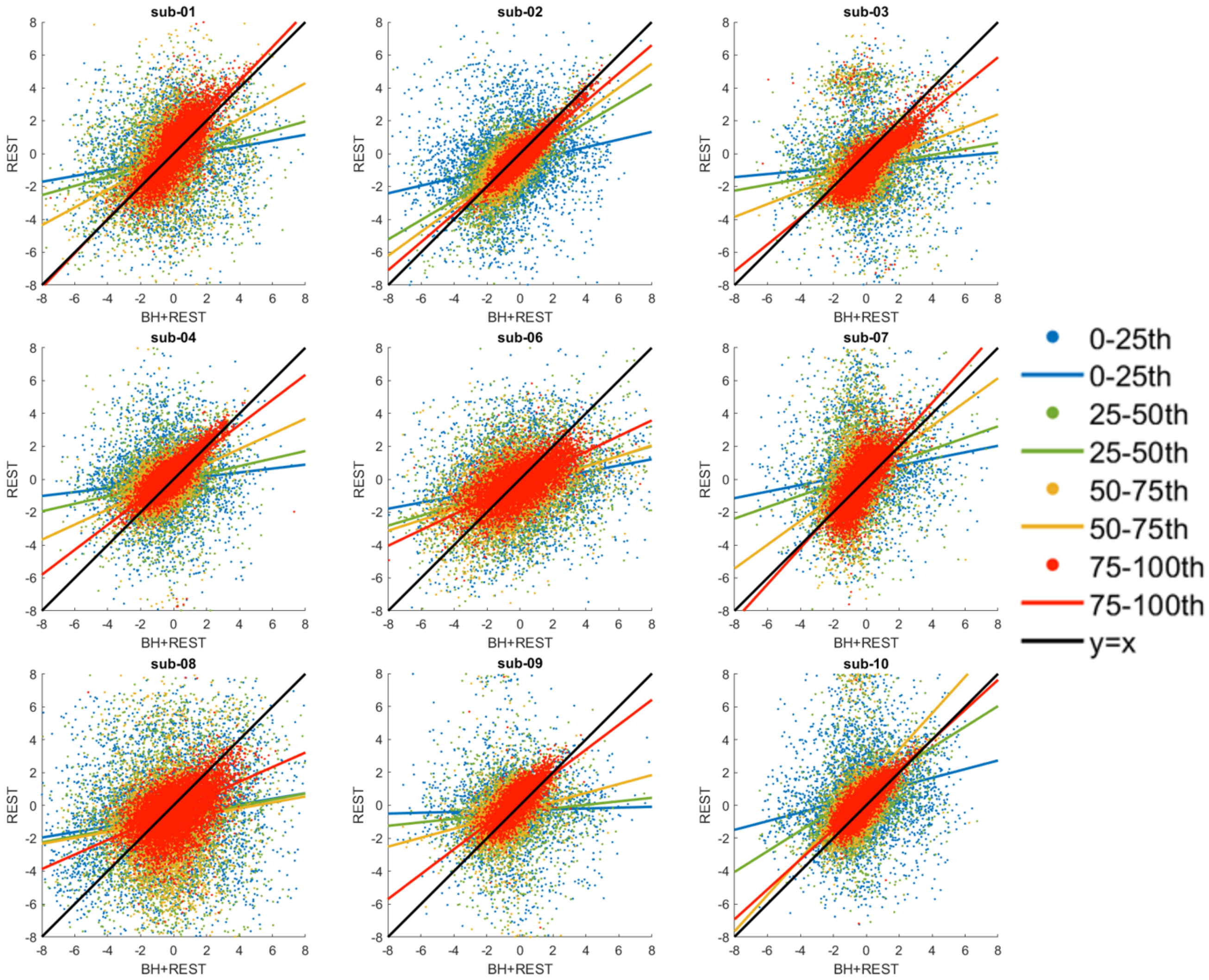


***Supplementary Figure 5.*** *Spatial correlation, in GM voxels, between hemodynamic delay times from the BH+REST and REST data segments, for all subjects. This is the same as what is shown in Supplementary Figure 2 (Panel A), but instead of filtering the voxels with a single amplitude threshold, amplitude percentile threshold bands are used (a voxel must be within the percentile threshold band to be included).*

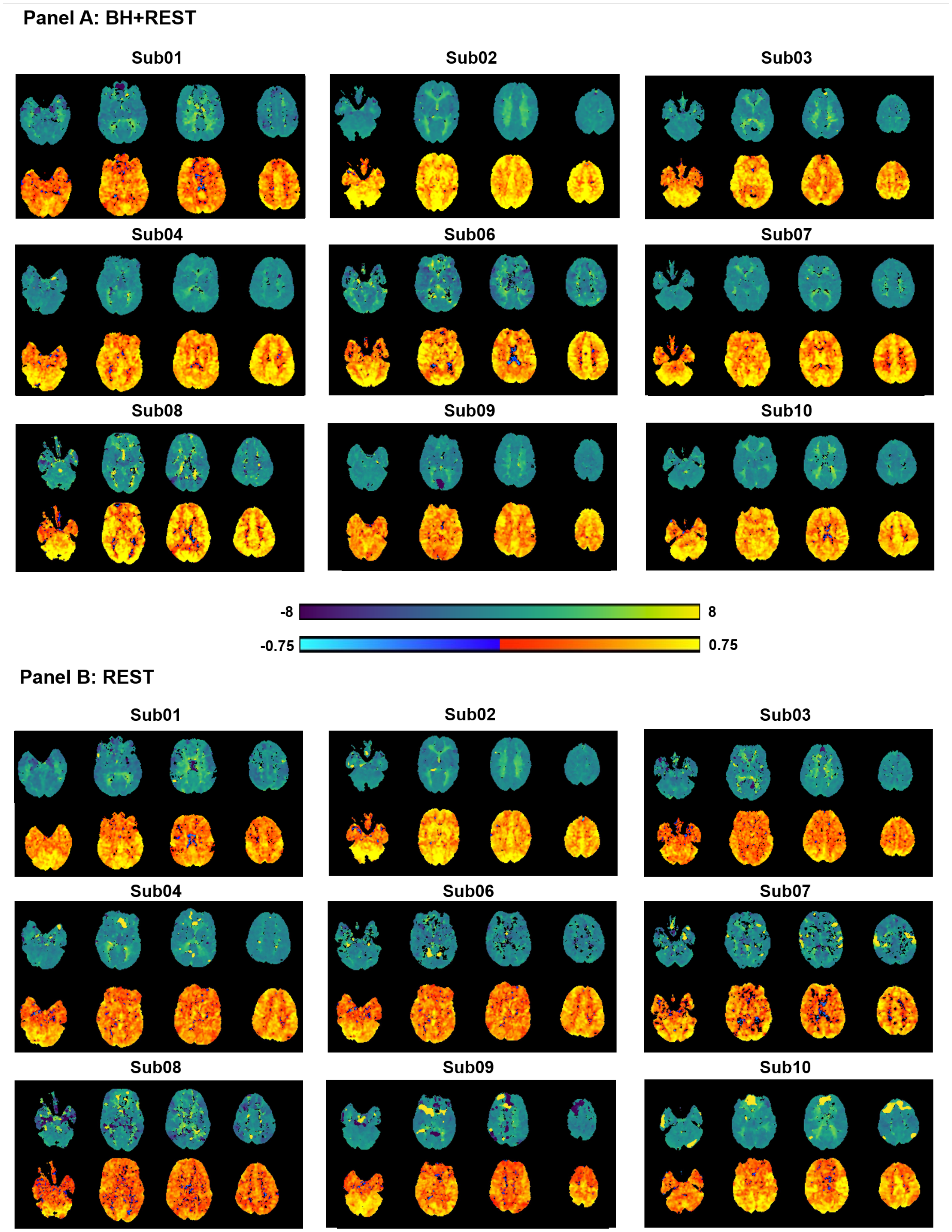


***Supplementary Figure 6.*** *Parameter maps generated with Rapidtide, for all example subjects, after increasing the level of smoothing (compared to Supplementary Figure 1).*

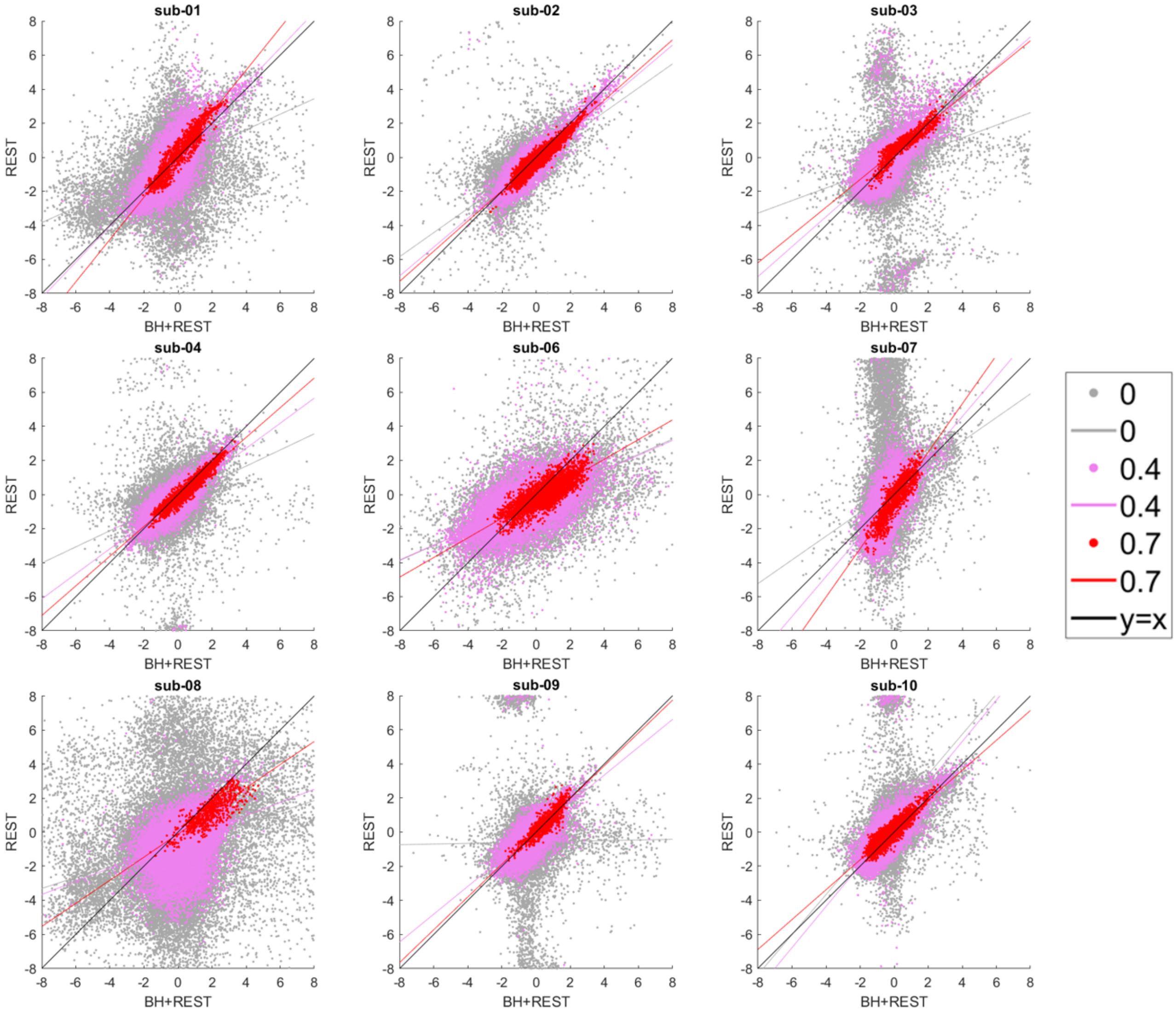


***Supplementary Figure 7.*** *Spatial correlation between hemodynamic delay times from the BH+REST and REST data segments, for all subjects, after increasing the level of smoothing (compared to Supplementary Figure 2). Results are grouped by amplitude threshold.*

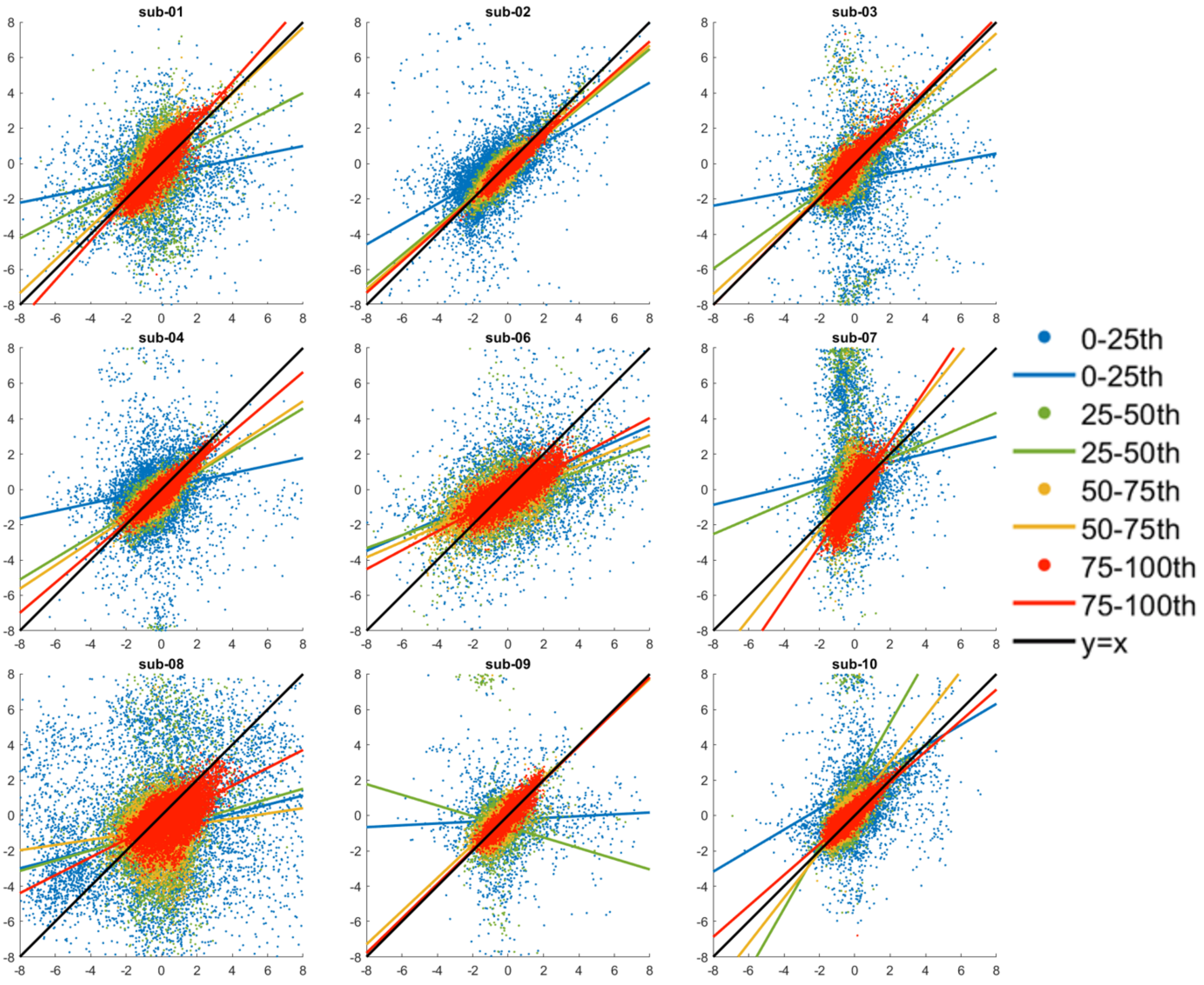


***Supplementary Figure 8.*** *Spatial correlation between hemodynamic delay times from the BH+REST and REST data segments, for all subjects, after increasing the level of smoothing (compared to Supplementary Figure 5). Results are grouped by amplitude percentile threshold band.*

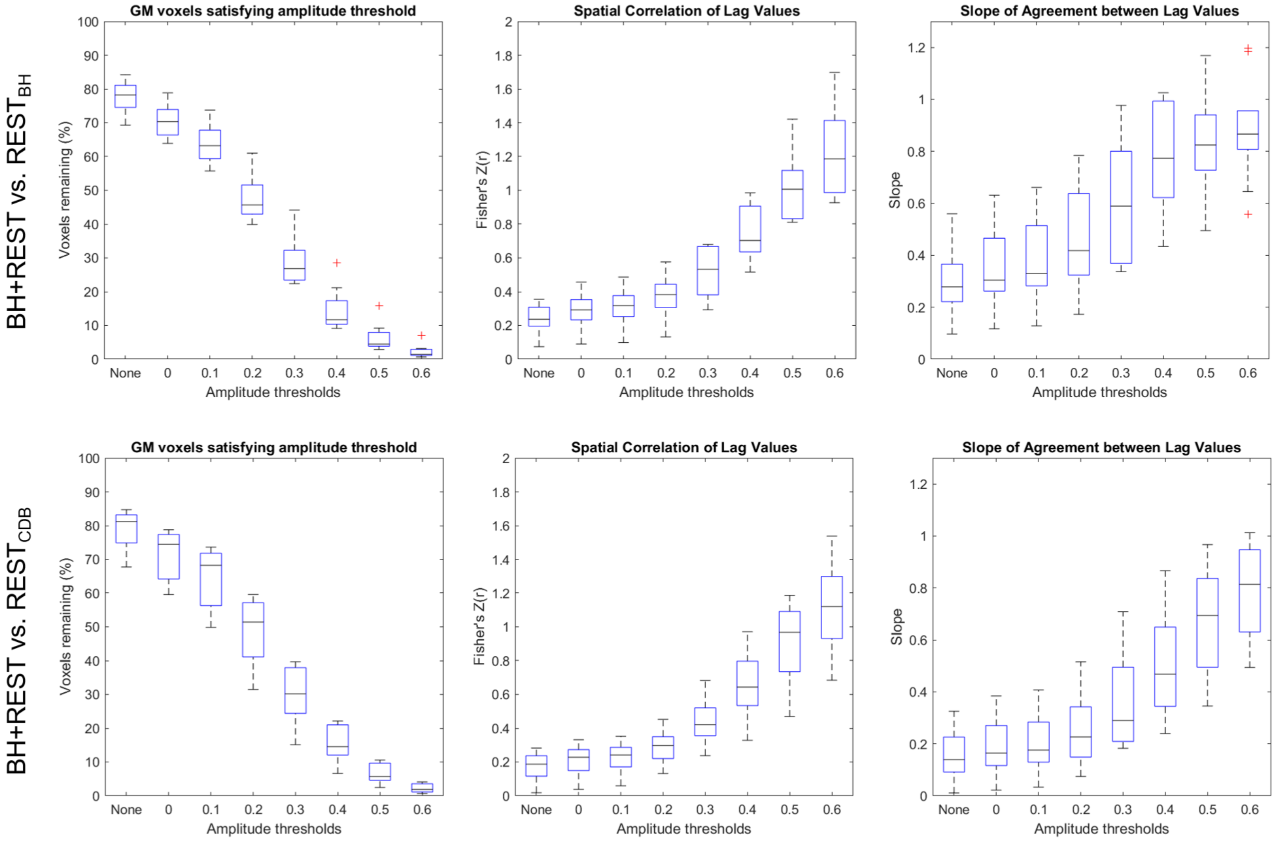


***Supplementary Figure 9.*** *Top row: same as Figure 2, comparing BH+REST and REST segments from the same acquisition (overlapping data); bottom row: comparing BH+REST and REST segments from different acquisitions (no overlap). Subject 6 was not included in the bottom comparison as they were previously deemed an outlier.*

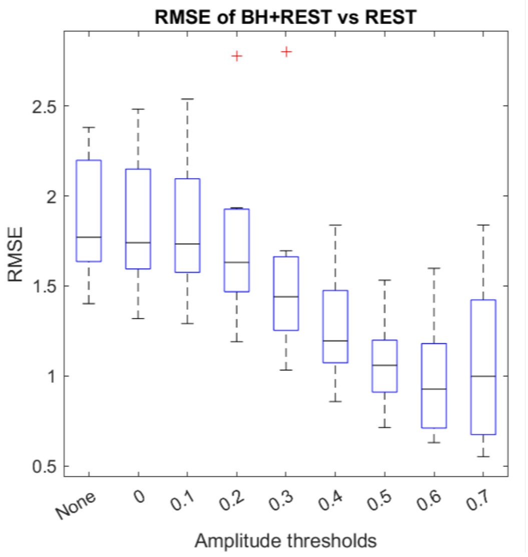


***Supplementary Figure 10.*** *Summary of the root mean square error (RMSE) of the linear fits at different amplitude thresholds (assessing the same relationship as displayed in Figure 2 but showing RMSE as an alternative to Pearson's r value).*

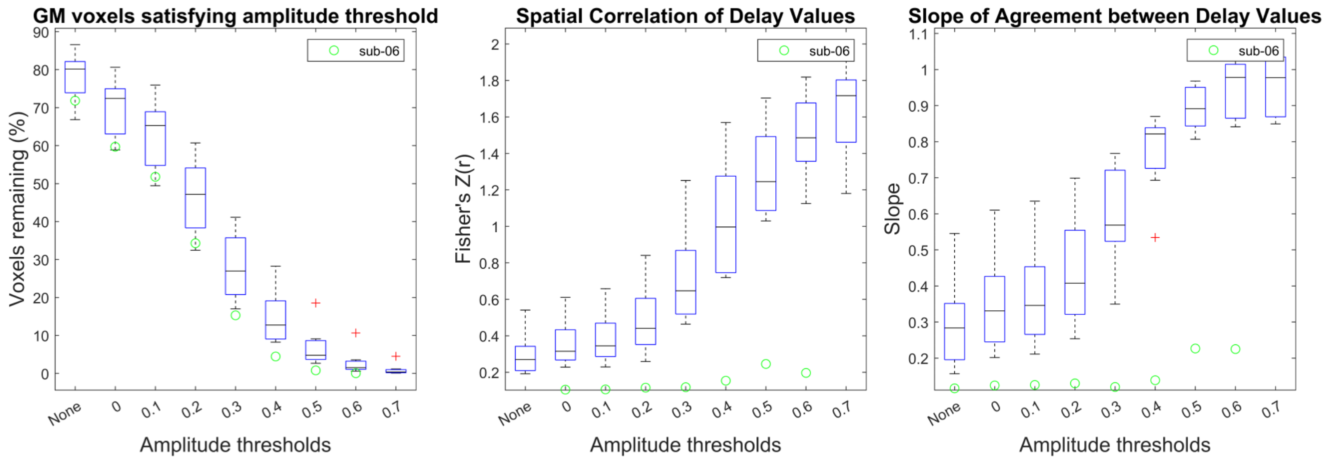


***Supplementary Figure 11.*** *Delay agreement between CDB+REST and REST data segments within GM voxels, for each amplitude threshold. The boxplots summarize across subjects. The left plot shows the percentage of GM voxels satisfying each amplitude threshold in both data segments, where 'None' refers to no amplitude thresholding applied, and the subsequent numbers (0 to 0.7) refer to the amplitude threshold the voxel must exceed in order to be included in the assessment of delay agreement. The middle and right plots summarize the delay agreement for each amplitude threshold, showing the spatial correlation coefficients (middle plot) and slopes (right plot). Sub-06 had too few voxels remaining after the 0.7 amplitude threshold; therefore, it was listed separately in the figures and only extends to the amplitude threshold of 0.6. This subject is also an outlier in spatial correlation values and slope, across other amplitude thresholds, compared to other subjects.*

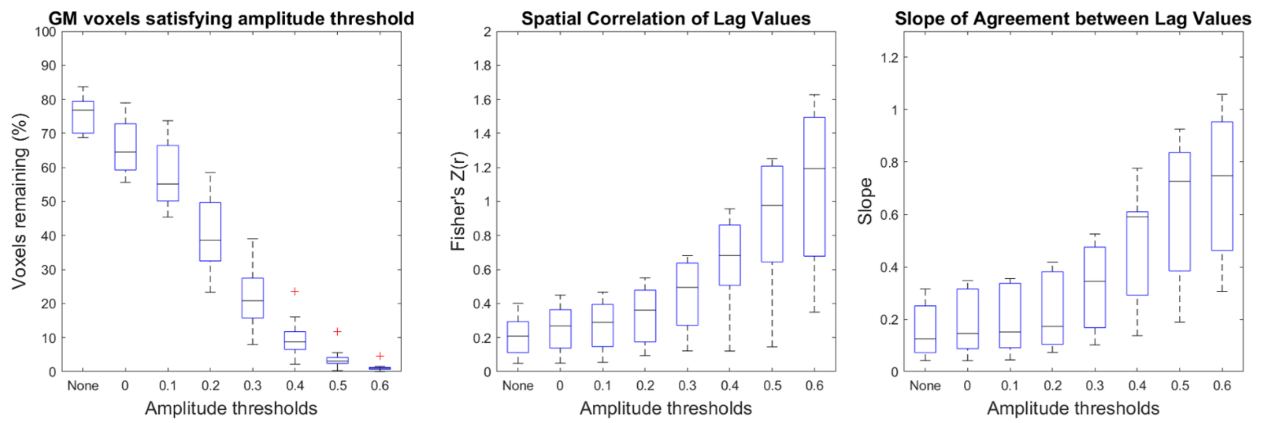


***Supplementary Figure 12.*** *Delay agreement between BH+REST and REST data segments within GM voxels, for each amplitude threshold using the Superior Sagittal Sinus reference time series. The boxplots summarize across subjects. The left plot shows the percentage of GM voxels satisfying each amplitude threshold in both data segments, where 'None' refers to no amplitude thresholding applied, and the subsequent numbers (0 to 0.6) refer to the amplitude threshold the voxel must exceed in order to be included in the assessment of delay agreement. The middle and right plots summarize the delay agreement for each amplitude threshold, showing the spatial correlation coefficients (middle plot) and slopes (right plot).*
